# A secreted endoribonuclease ENDU-2 from the soma protects germline immortality in *C. elegans*

**DOI:** 10.1101/2020.12.04.408260

**Authors:** Wenjing Qi, Erika D v. Gromoff, Fan Xu, Qian Zhao, Wei Yang, Dietmar Pfeifer, Wolfgang Maier, Lijiang Long, Ralf Baumeister

## Abstract

Multicellular organisms coordinate tissue specific response to environmental information via both cell-autonomous and non-autonomous mechanisms. In addition to secreted ligands, secreted small RNAs have recently been reported to regulate gene expression across tissue boundaries. Here we show that the conserved poly-U specific endoribonuclease ENDU-2 is secreted from the soma and taken-up by the germline to ensure germline immortality at elevated temperature in *C. elegans*. ENDU-2 binds to mature mRNAs and negatively regulates mRNA abundance both in the soma and the germline. While ENDU-2 promotes RNA decay in the soma directly via its endoribonuclease activity, ENDU-2 prevents misexpression of soma-specific genes in the germline and preserves germline immortality independent of its RNA-cleavage activity. In summary, our results suggest that the secreted RNase ENDU-2 transmits environmental information across tissue boundaries and contributes to maintenance of stem cell immortality probably via retaining a stem cell specific program of gene expression.

## Introduction

Germ cells are the only type of cells within an organism that deliver genetic and epigenetic material to the offspring. Germ cells are therefore distinct from the somatic cells in their ability to maintain totipotency to produce an entire organism upon fertilization and their immortality to allow reproduction for unlimited future generations. In *C. elegans*, loss of germline immortality typically results in sterility of initially fertile animals after a number of generations. To date, telomerase-mediated telomere maintenance, balanced heterochromatic H3K9 methylation and euchromatic H3K4/H3K36 methylation are the prevalent known molecular mechanisms involved in maintenance of germline immortality (Ahmed and Hodgkin, 2000; Katz et al., 2009; Lev et al., 2017; Weiser et al., 2017). Piwi RNA mutants and mutations in multiple components of a nuclear RNA interference (RNAi) pathway that promotes the inheritance of germline RNAi also result in a temperature-dependent mortal germline (Mrt) phenotype (Buckley et al., 2012; Simon et al., 2014). Failure to maintain germline immortality in Piwi RNA mutants has recently been explained by transgenerational silencing of histone genes (Barucci et al., 2020). These studies, however, focused almost exclusively on autonomous mechanisms in the germline. It is unknown whether somatic signaling may contribute to maintenance of germline immortality.

*C. elegans* ENDU-2 belongs to a conserved but less studied family of proteins containing proposed poly-U specific endoribonuclease (XendoU) domains. While human EndoU (PP11 placental protein 11) is used as a cancer marker gene, Viral EndoU Nsp15 is highly conserved in all known coronavirus and has been suggested to promote viral RNA replication and limit innate immunity response of the host cells (Hackbart et al., 2020; Ivanov et al., 2004). *C. elegans* ENDU-2 has been shown to regulate cold stress response (Ujisawa et al., 2018). Very recently, ENDU-2 is reported to regulate nucleotides metabolism and germline proliferation in response to changes in nucleotide levels and genotoxic stresses (Jia et al., 2020).

Here we report that secretion of the poly-U specific endoribonuclease ENDU-2 from the soma to the germline preserves germline immortality at elevated temperature. We find that ENDU-2 binds to mature mRNAs and down-regulates mRNA levels both in the soma and the germline. In addition, RNA-binding and -cleavage are two separable activities utilized by ENDU-2 to control gene expression via distinct mechanisms. In the soma, ENDU-2 relies on its endoribonuclease activity to negatively regulate part of its mRNA targets. In the germline, ENDU-2 prevents misexpression of soma-specific genes and ensures stem cell immortality primarily only via its RNA-binding activity, suggesting an essential role of ENDU-2 in retaining the stem cell specific program of gene expression. In summary, our data implicates that the soma sends ENDU-2 as a messenger to the germline to control gene expression across tissue boundaries in response to temperature alteration.

### Loss of *endu-2* causes a temperature dependent Mrt phenotype

Freshly outcrossed *endu-2(lf)* mutants were similar to wild type animals in the first few generations, except slightly reduced germline proliferation, an egg-laying defect (Egl) due to abnormal development of the vulva, and a moderate reduction in adult lifespan at 20°C (Supplemental Fig. S1). Long-term strain maintenance at 20°C was difficult, since the *endu-2(lf)* mutants showed gradually increased sterility that, however, could be reset by new outcrosses with wild type animals. We examined the fertility of two *endu-2* alleles *tm4977* and *by188* over generations after four additional out-crosses. *tm4977* allele has a 620 bp deletion starting from the 5’UTR to the end of the third intron, while the 20 bp deletion in *by188* causes an early stop codon in the first exon. Therefore, these two alleles are probably null mutants. Both *tm4977* and *by188* alleles became sterile at 20°C after about 15-20 generations (Fig. 1A, Supplemental Fig. S2A). Since we did not observe a significant reduction of the number of germ nuclei in the mitotic zone across generations, sterility was not the consequence of declining germline proliferation (Supplemental Fig. S2B). Instead, *endu-2* day 1 adult animals in the generations with highly penetrant sterile phenotype showed pleiotropic defects in the germline. The most prominent defects were abnormal cell death (38%, n=105) and increased number of apoptotic corpses (31% with ≥ 2 corpses per gonad arm, n=105) (Fig. 1B). We also observed ongoing spermatogenesis 24 hours after mid-L4 stage (11%, n=105) whereas wild type animals had completely switched spermatogenesis to oogenesis, suggesting prolonged spermatogenesis. Furthermore, endomitosis occurred in the mitotic region (14%, n=25) (Fig. 1B). In the generations displaying strong sterile phenotype, *endu-2(lf)* additionally showed high incidence of male (Him, 14%, n=218) and increased occurrence of other phenotype aspects (Rol, Dpy, Sma phenotypes, in total 10%, n=218). The latter was probably caused by elevated somatic mutation rates, since these phenotypes were not heritable. The sterile phenotype was 100% penetrant at 25°C already at generation 6-10, but was not detected at 15°C (Supplemental Fig. S2C and S2D). Notably, the highly penetrate sterility could be fully reversed within several generations by transferring the animals to 15°C (Fig. 1A). Taken together, these results suggest an essential role of ENDU-2 in preserving germline immortality at elevated temperature and existence of additional mechanisms to compensate for the loss of *endu-2* at 15°C.

**Fig. 1.**
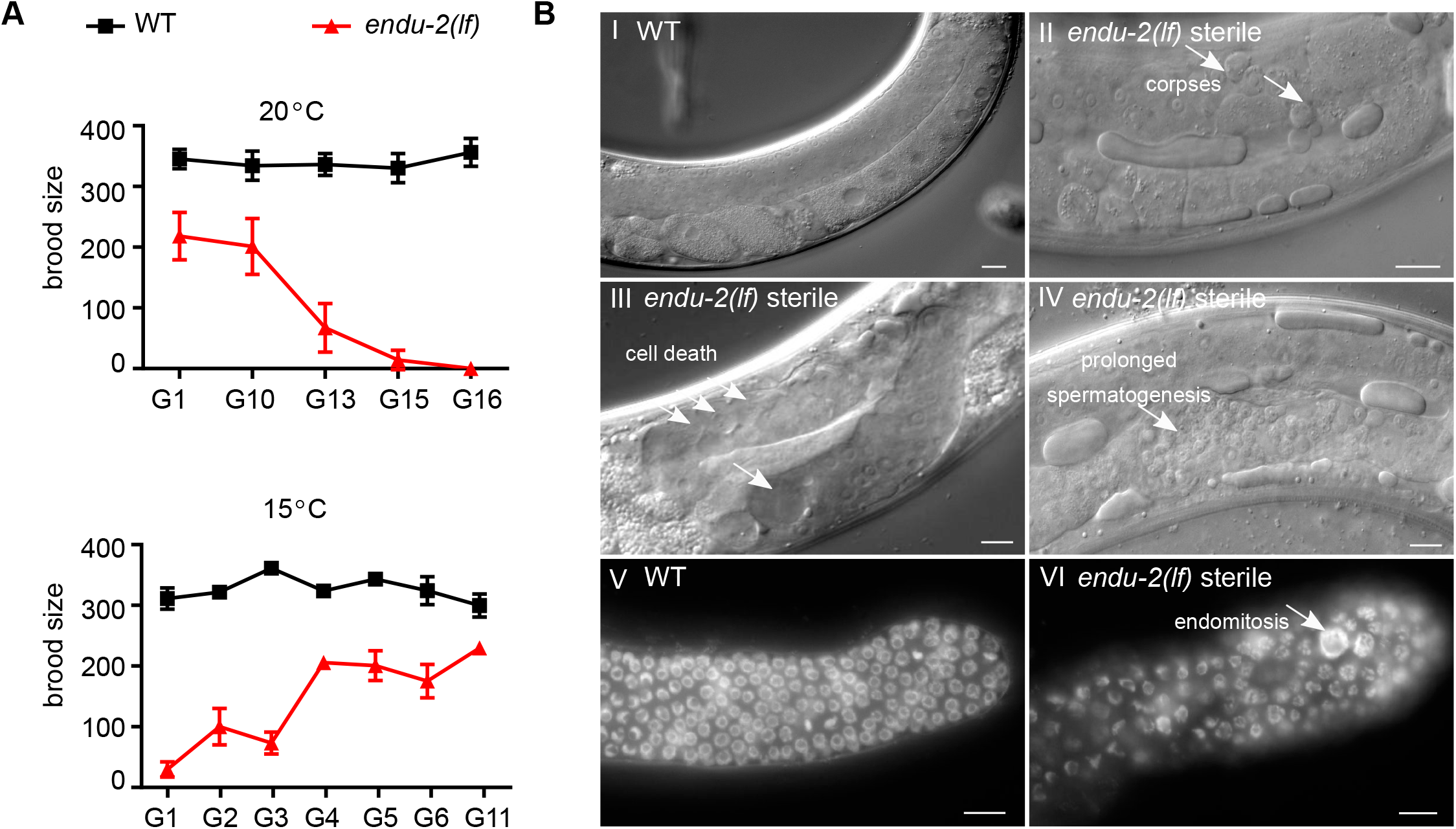
*endu-2* mutant shows a temperature dependent Mrt phenotype. **A**. Upper graph: Brood size of *endu-2(tm4977)* animals decreases gradually after 4x initial backcrosses with wild type (G0) over generations at 20°C. Lower graph: The defects of *endu-2(tm4977)* animals in reproduction is reversible at 15°C. To obtain this graph, adult animals from G6 that had been maintained at 25°C were shifted back to 15°C and counted as G0. As numbers of generation to reach 100% sterility at 20°C among the 4 biological replicates varied from 10 to 20, only one represented replicate with n=15 animals for each generation is shown. The data are mean ± SEM. **B**. The sterile *endu-2(tm4977)* animals display multiple defects in germline mitosis and meiosis. I-IV) DIC images of wild type and sterile *endu-2(tm4977)* animals 24 hours after mid-L4 stage. The white arrows point to (II) increased number of germline apoptotic corpses; (III) empty gonad due to germ cell death; IV) prolonged spermatogenesis; (V) DAPI staining of germline proliferating zone of day one wild type and (VI) sterile *endu-2(tm4977)* adults, respectively. White arrows in VI point to abnormally large chromosomes due to endomitosis. Scale bar 10 μm.

### ENDU-2 is a secreted protein

A previous report has suggested a wide-spread expression of *endu-2* in somatic tissues (Ujisawa et al., 2018), but did not report ENDU-2 localization in the germline. To retest the expression pattern of ENDU-2, we generated several independent transcriptional, translational EGFP fusion *endu-2* reporters and a CRISPR/Cas9 EGFP knock-in strain at the endogenous *endu-2* locus (expression constructs are shown in the Supplemental Fig. S3A). The transcriptional fusion reporter gene, harboring 4 kb of *endu-2* upstream sequences, was expressed only in the intestine (Supplemental Fig. S3B). A translational reporter harboring the same upstream sequences, and also the entire genomic region of *endu-2*, fused to EGFP, rescued Egl, reduced germline proliferation, and short lifespan phenotypes (Supplemental Fig. S1A, S1C and S1D). The CRISPR/Cas9 EGFP knock-in strain displayed neither Mrt nor Egl phenotypes (Supplemental Fig. S1E). These results suggest that ENDU-2::EGFP fusion proteins in the transgenic and the EGFP knock-in strains are functional. We observed ENDU-2::EGFP protein in the cytoplasm of intestine, somatic gonad and coelomocytes in both *endu-2::EGFP* transgenic and the CRISPR/Cas9 EGFP knock-in strains (Fig. 2A and Supplemental Fig. S3C). Different from the study of Ujisawa et al. (2018), we did not detect ENDU-2::EGFP in the musculature or in the neurons. Instead, we noticed extracellularly localized ENDU-2::EGFP, such as in the interspace between uterine wall and embryos (Fig. 2A and Supplemental Fig. S3C), suggesting that ENDU-2 may be a secreted protein. ENDU-2 indeed contains a predicted N-terminal (1-19 amino acid) secretion signal peptide for endoplasmic reticulum (ER) targeting, indicating that the protein is destined towards the secretory pathway. Fusion of this secretion signal peptide to the N-terminus of a neuron specific expressed EGFP reporter led to strongly decreased expression level of EGFP in neurons accompanied by EGFP signal in extracellular space and coelomocytes, which endocytose secreted proteins existing in the pseudocoelic fluid (Fig. 2B). In addition, removing the N-terminal secretion signal peptide of ENDU-2 (Δ_ss_ENDU-2::EGFP) resulted in strong expression of Δ_ss_ENDU-2::EGFP only in the intestine but not in the coelomocytes (Supplemental Fig. S3E and S4A). These results together suggest that the secretion signal peptide composed of the first 19 amino acids of ENDU-2 is necessary and sufficient to trigger secretion of a protein. Strikingly, we noticed that another transgenic reporter expressing 3xFlag::ENDU-2::EGFP was expressed strongly in the intestinal and also weakly in some head neurons, muscle cells in the head region and anal depressor muscle cells, corroborating the report from Ujisawa et.al (Supplemental Fig. S3D). We speculate that the N-terminal 3xFlag fusion may prevent secretion by impairing binding of the secretion signal peptide by signal recognition particle (SRP), thus allowing detection of the weak expression in the muscle and neuron. In addition, expression in the head neurons and muscle cells might possibly be controlled by promoter signals localized in the first intron since this was the only sequence absent in the transgene expressing Δ_*ss*_*endu-2::EGFP* (Supplemental Fig. S3A), in which we failed to observe muscle or neuronal localization. Moreover, when we expressed *endu-2::EGFP* selectively either in the neurons (*unc-119* promoter), muscles (*myo-3* promoter) or intestine (*vha-6* promoter), ENDU-2::EGFP was always detected in the coelomocytes (Supplemental Fig. S4A), indicative for its secretion from these tissues. Notably, expressing *endu-2::EGFP* specifically in either neurons or muscles of heat-stressed animals resulted in ENDU-2::EGFP localization in the pharynx (Supplemental Fig. S4B), suggesting that either secretion or uptake of ENDU-2 in these tissues could be modulated by temperature.

**Fig. 2.**
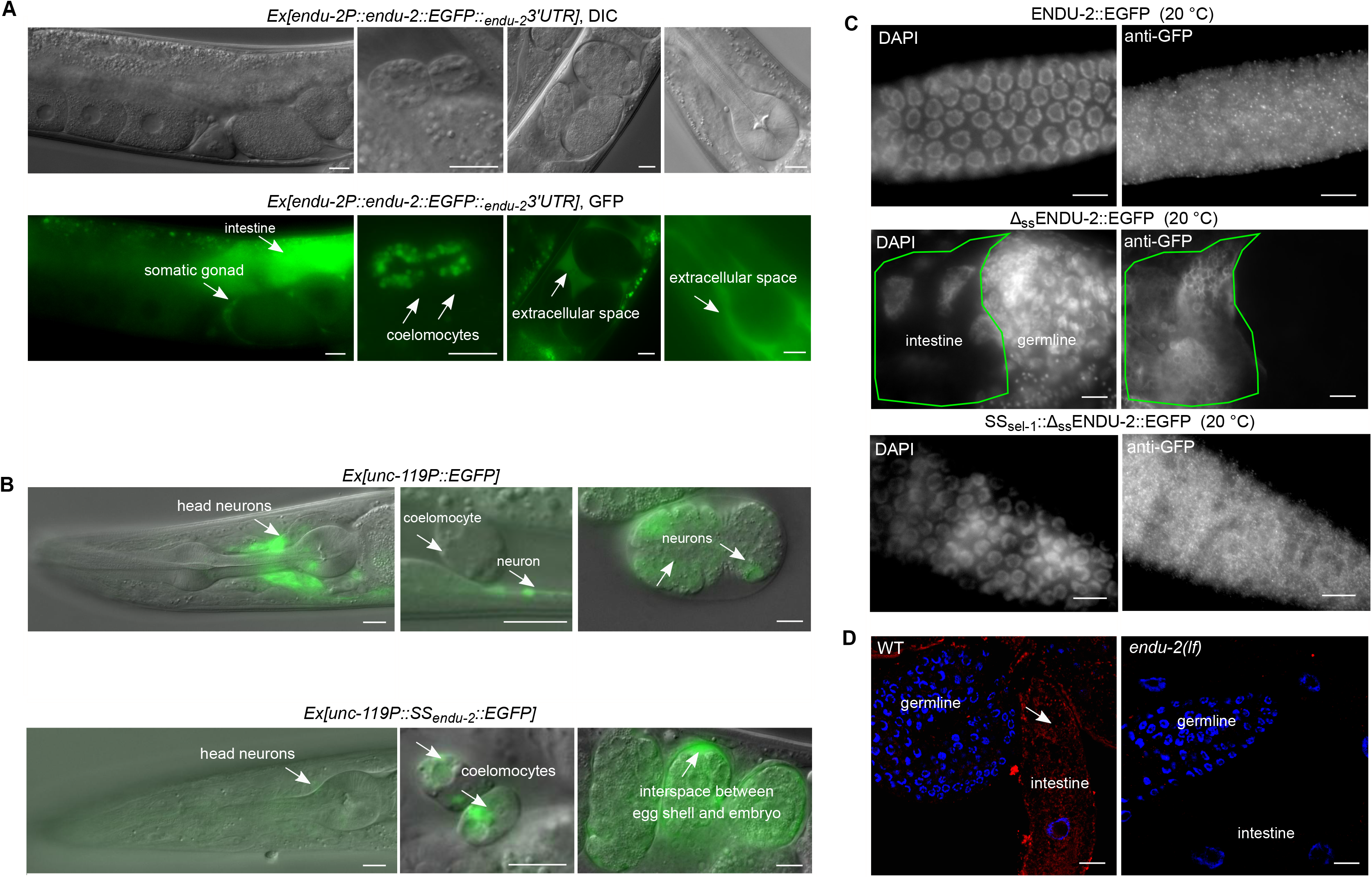
ENDU-2 is a secreted protein. **A.** Fluorescence micrographs of transgenic *endu-2(tm4977);byEx1814[endu-2P::endu-2::EGFP::_endu-2_3’UTR]* animals. ENDU-2::EGFP is detected in the intestine, the somatic gonad, coelomocytes and extracellular space between the uterus and embryos. **B.** Fusion of the 1-19 amino acids of ENDU-2 (SS_endu-2_) to neuronal specific expressed EGFP is sufficient for secretion of EGFP protein. Upper panels shows expression pattern of *unc-119P::EGFP* which is detected only in neuronal cells (strong) of animals and embryos. Lower panels are images showing expression pattern of *unc-119P::SS_endu-2_::EGFP* which is detected in the neurons (weak), coelomocytes and interspace between egg shell and embryos, indicative of efficient secretion of EGFP enforced by the secretion signal peptide of ENDU-2. Images are representative for more than 20 animals, analyzed by using a 40× objective. **C.** Fluorescence micrographs of GFP-antibody staining of *endu-2(tm4977);byEx1375[endu-2P::endu-2::EGFP]*, *endu-2(tm4977);byEx1449[endu-2P::Δ_ss_endu-2::EGFP]* and *endu-2(tm4977);byEx1875[endu-2P::SS_sel-1_::Δ_ss_endu-2::EGFP]* transgenic animals at 20°C. ENDU-2::EGFP (n=62, 100%) and SS_sel-1_::Δ_ss_ENDU-2::EGFP (n=21, 100%) but not Δ_ss_ENDU-2::EGFP (n=26, 0%) is detected in the germline. N=2. **D.** smFISH staining reveals presence of *endu-2* mRNA in the intestine (white arrow) but not in the germline. Blue: DAPI stained nuclear DNA. Red: smFISH probes stained mRNA. N=4. Scale bar 10 μm for all images in this Figure.

### Secretion of ENDU-2 from the soma to the gonad protects germline immortality

It is generally accepted that multi-copy extrachromosomal arrays are silenced and not expressed in the *C. elegans* germline (Kelly et al., 1997). However, we found that extrachromosomal *endu-2::EGFP* transgenes rescued the Mrt germline phenotype of *endu-2(lf)* animals (Fig. 3A), suggesting that somatic ENDU-2 might preserve germline immortality across tissue boundaries. Our hypothesis was that secreted ENDU-2 could be endocytosed by the gonad. To test this assumption, we first tested whether ENDU-2 protein could be detected in the germline. All of our *endu-2::EGFP* transgenic reporters displayed weak expression level unless secretion was blocked. Therefore, we could only occasionally observe faint ENDU-2::EGFP in few oocytes (Supplemental Fig. S4C). Via GFP antibody staining we detected ENDU-2::EGFP in punctate structures in the germline at 15°C, 20°C and 25°C (Fig. 2C and Supplemental Fig. S5). In contrast, Δ_ss_ENDU-2::EGFP was not detectable in the germline despite of its high expression level (Supplemental Fig. S3F). And fusion the secretion signal peptide of another secreted protein SEL-1 (Grant and Greenwald, 1996) to Δ_ss_ENDU-2::EGFP reactivated secretion and its localization in the germline (Fig. 2C and Supplemental Fig. S3E). These observations indicate that secretion of ENDU-2 is indispensable for its gonadal localization. Moreover, we performed single molecular FISH (smFISH) staining to visualize *endu-2* mRNA. *endu-2* mRNA signals were visible predominately in the intestine but clearly absent in the germline (Fig. 2D). Furthermore, sequencing of RNA extracted from isolated gonads also revealed absence of *endu-2* mRNA in the wild type germline (Supplemental Table S3). Taken together, these results implicate that somatically expressed ENDU-2 protein can enter the gonad via its secretion and uptake.

**Fig. 3.**
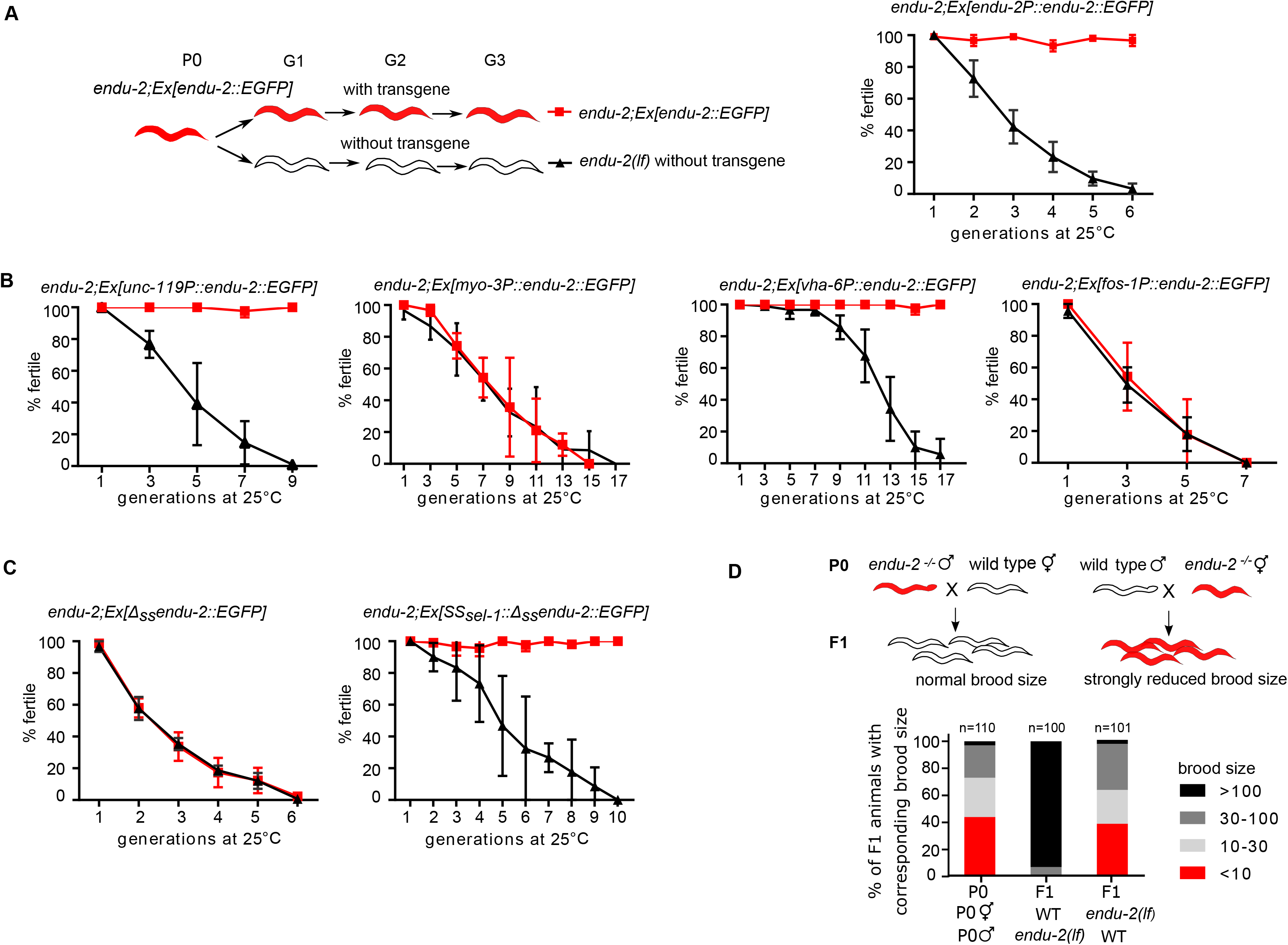
Maternally provided ENDU-2 from the soma contributes to maintenance of germline immortality. **A**. *endu-2(wt)::EGFP* rescues the Mrt phenotype at 25°C. Left panel shows the experimental strategy for all rescue experiments with *endu-2* transgenes in this study. *endu-2(−/−)* daughter generation (G1) that has lost the extrachromosomal rescue constructs expressing *endu-2::EGFP* was isolated. This and the following *endu-2(−/−)* generations (G2 to Gn) were compared to *endu-2;Ex[endu-2:.EGFP]* derived from the same P0 animal. Data are mean ± SD, N=3. **B**. Expression of *endu-2(wt)::EGFP* from neuron or intestine is sufficient to rescue the Mrt phenotype at 25°C. Data are mean ± SD, N=3. **C**. *SS_sel-1_::Δ_ss_endu-2::EGFP* but not *Δ_ss_endu-2::EGFP* transgene rescues the Mrt phenotype at 25°C. Data are mean ± SD, N=3. **D**. ENDU-2 primarily affects oocyte to maintain germline immortality. Data are pooled data from three biological replicates with similar tendencies in results.

Next, we asked from which tissue ENDU-2 ensures germline immortality. Expressing ENDU-2 specifically in the intestine or the neurons, but not the muscle or somatic gonad, were sufficient to rescue the Mrt phenotype, indicating a non-cell-autonomous ENDU-2 signal from the neurons and intestine to the germline (Fig. 3B). In addition, expressing the secretion deficient Δ_ss_ENDU-2::EGFP failed to rescue the Mrt phenotype (Fig. 3C). Furthermore, SS_sel-1_∷ Δ_ss_ENDU-2::EGFP rescued the Mrt phenotype (Fig. 3C), suggesting that guiding ENDU-2 into ER-Golgi secretory pathway rather than the first 19 amino acids of ENDU-2 is essential for its function in the germline.

To directly test whether loss of somatic *endu-2* expression is sufficient to induce a mortal germline, we performed *endu-2* RNAi knock-down in a *ppw-1* mutant background in which germline RNAi does not function (Tijsterman et al., 2002). Although neither wild type nor *ppw-1* mutants upon *endu-2* RNAi displayed a fully penetrant Mrt phenotype within 15 generation, *endu-2* RNAi, resulted in gradually reduced brood sizes over generations in both *ppw-1* and wild type background, indicating that somatically expressed *endu-2* mRNA is required for normal reproduction (Supplemental Fig. S6A). The failure to obtain a fully penetrant Mrt phenotype in this experiment may be due to the relative inefficiency of RNAi in the nervous system that, expresses *endu-2* to ensure germline immortality.

### ENDU-2 affects the oocytes to preserve germline immortality independent of the nuclear RNAi pathway

To know whether ENDU-2 affects oocytes or sperm to ensure germline immortality, we tested the parental contribution of ENDU-2 during sexual reproduction. For this purpose, we crossed *endu-2(*+/+) wild type parents with either males or hermaphrodites of *endu-2(−/−)* that had been grown at 25°C for five to seven generations and displayed strongly reduced brood size and high percentage of sterile phenotype (Fig. 3D). Whereas the heterozygous progeny of *endu-2(*+/+) mothers had brood sizes with typically more than 100 F2 animals, the brood size of F1 cross-progeny derived from *endu-2(−/−)* mothers was similarly low as that of their mothers. We conclude that ENDU-2 function is required predominantly in the oocytes to preserve germline immortality.

The temperature dependent Mrt phenotype in *endu-2* mutants resembles that of *hrde-1* mutants in the nuclear RNAi pathway (Buckley et al., 2012; Ni et al., 2016). In addition to loss of germline immortality, *hrde-1* is also defective in multigenerational inheritance of germline RNAi. To know whether ENDU-2 acts in the nuclear RNAi pathway, we examined *oma-1* RNAi inheritance, which leads to suppression of the embryonic lethality of the *oma-1* gain-of-function mutant for several generations (Alcazar et al., 2008). Unlike the *hrde-1* mutants that lost inheritance of *oma-1* RNAi within two generations, effect of *oma-1* RNAi knock-down persisted for 5-6 generations both in *endu-2(lf)* and wild type animals (Supplemental Fig. S6B), suggesting that ENDU-2 does not act in the nuclear RNAi pathway to control transgenerational inheritance of germline RNAi. Therefore, maintenance of germline immortality by ENDU-2 probably functions via a distinct mechanism from that of the nuclear RNAi pathway.

### mRNA-binding, but not mRNA-cleavage, by ENDU-2 is essential for maintaining germline immortality

ENDU-2 harbors two XendoU domains, of which the C-terminally localized one is more similar to both human EndoU and *Xenopus* XendoU (Supplemental Fig. S7). It had been suggested recently that ENDU-2 might also be an RNA-binding protein (Ujisawa et al., 2018). To identify the candidate RNAs associated with ENDU-2, we precipitated ENDU-2::EGFP and analyzed co-immunoprecipitated RNAs by deep-sequencing (RIP-Seq). We also reasoned that, since wild type ENDU-2 would potentially cleave its RNA targets, this might prevent enrichment of intact RNA targets for identification. Although we had no direct proof of an RNA cleaving activity of ENDU-2 yet, we thought that generation of an ENDU-2 variant that maintains RNA-binding, but loses cleavage activity, should facilitate RNA target detection. *Xenopus* XendoU mutants with E to Q exchanges at positions 161 or 167 of the EndoU domain lose RNA-cleavage without reducing RNA-binding activity (Gioia et al., 2005). The second glutamic acid (167E) is conserved in both XendoU domains of *C. elegans* ENDU-2 (175E and 460E), whereas the first (161E) is only found in the second XendoU domain (454E) (Supplemental Fig. S7B). We expressed both ENDU-2(E454Q)::EGFP and ENDU-2(E460Q)::EGFP variants in *endu-2(lf)* background and performed RIP-Seq with these two strains in additional to wild type ENDU-2::EGFP. By plotting normalized reads (RPM) of each transcript we were able to investigate RNA binding affinity of ENDU-2 under different conditions. In general, all three ENDU-2 variants tested showed stronger RNA-binding affinity at 15°C than at 25°C (Fig. 4A). E460Q showed weaker RNA binding already at 15°C than wild type ENDU-2 and almost completely lost RNA-binding capacity at 25°C. In addition, ENDU-2(E454Q) displayed stronger RNA binding activity than ENDU-2(wt) at 15°C. Therefore, we used RIP-Seq data of ENDU-2(E454Q) at 15°C for detecting RNAs bound by ENDU-2. A total of 5,920 transcripts were co-immunoprecipitated with ENDU-2(E454Q) (Supplemental Table S1). Most of them (>99%) were protein coding transcripts except few non-coding RNAs (5 snoRNAs, 27 pseudogenes, 5 ncRNAs and 1 lincRNA). In addition, reads distribution analysis showed > 99% of sequenced reads were mapped to exonic regions, suggesting that ENDU-2(E454Q) primarily bound to processed mRNAs (Fig. 4B). We performed *in vitro* mRNA binding assays with recombinant ENDU-2 proteins and two selected mRNA targets from the RIP-Seq data and could confirm that both ENDU-2 and ENDU-2(E454Q) directly interacts with these mRNAs (Fig. 4C).

**Fig. 4.**
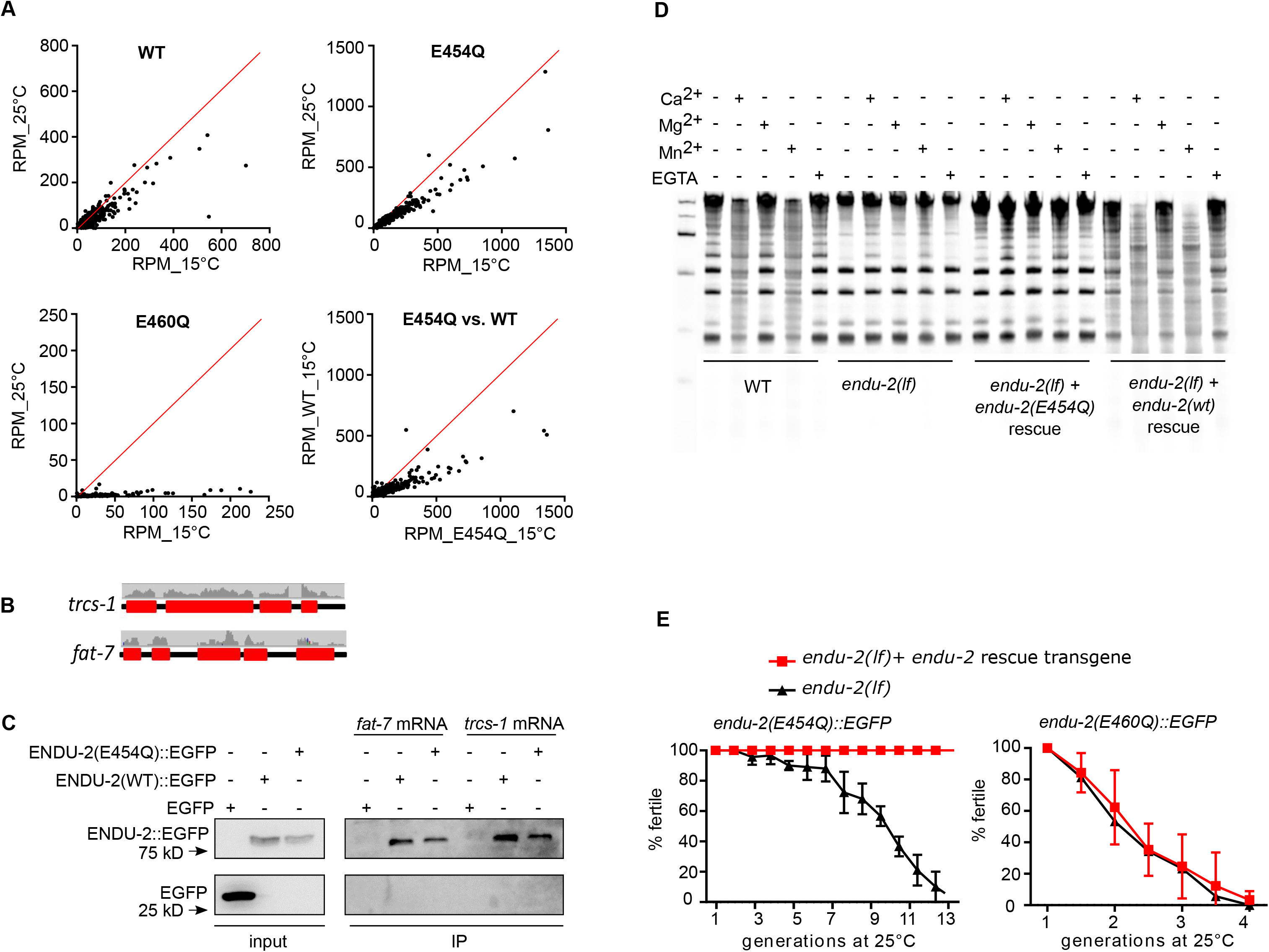
mRNAs-binding but not -cleavage activity of ENDU-2 protects germline immortality. **A.** Comparison of RNA binding activities of ENDU-2(wt)::EGFP, ENDU-2(E454Q)::EGFP, and ENDU-2(E460Q)::EGFP at 15°C and 25°C. Shown are plotted normalized reads (RPM) from RIP-Seq of each identified transcript under different conditions. **B.** Mapping of the RIP-Seq reads of two representative binding targets of ENDU-2. Only fragments of mRNA exons (red boxes) but not introns (black lines) were co-immunoprecipitated with ENDU-2. **C.** ENDU-2::EGFP variants bind to selected mRNAs in *vitro*. Shown are Western Blots to detect proteins binding to *fat-7* and *trcs-1* mRNA, respectively. EGFP is a negative control. N= 3 biological replicates. **D.** Wild type ENDU-2, but not ENDU-2(E454Q), leads to RNA decay (smear) in a Ca^2+^ and Mn^2+^ dependent manner, N=3. Shown is fused images of two gels from one experiment. The results of additional two biological replicates are shown in the Supplemental Fig. S8. **E.** *endu-2(E454Q)::EGFP* but not *endu-2(E460Q)::EGFP* transgene rescues the Mrt phenotype. Data are mean ± SD, N=3. Both e*ndu-2(tm4977)* and *endu-2(tm4977)* carrying *endu-2(E454Q)::EGFP* or *endu-2(E460Q)::EGFP* transgenes were decedents of one single P0 animal carrying the respective transgenes.

Studies of *Xenopus* XendoU have shown that that the RNA hydrolysis activity of XendoU requires Mn^2+^ or Ca^2+^ (Schwarz and Blower, 2014). ENDU-2 seemed to have a similar requirement, since the addition of 5 mM Mn^2+^ or Ca^2+^ in the buffer medium, but not 5 mM Mg^2+^ led to degradation of bulk RNA in the wild type worm lysates (Fig. 4D and Supplemental Fig. S8). Bulk RNA degradation was strongly reduced in *endu-2(lf)* animals. In addition, expression of the wild type *endu-2::EGFP* transgene, but not *endu-2(E454Q)::EGFP*, restored RNA decay in *endu-2(lf)* mutants, supporting our hypothesis that ENDU-2(E454Q) lost RNA-cleavage capability despite of the increased RNA-binding. Surprisingly, extrachromosomal expression of ENDU-2(E454Q), but not of ENDU-2(E460Q), fully rescued the Mrt phenotype of *endu-2(lf)* at 25°C (Fig. 4E). We conclude that mRNA-binding, rather than -cleavage activity of ENDU-2, is essential in the germline to maintain stem cell immortality at elevated temperature.

### ENDU-2 negatively regulates somatic mRNA abundance via its endoribonuclease activity

To test if ENDU-2 affects mRNA level, we performed microarray experiments at 25°C to determine transcriptomic alterations mediated by ENDU-2 (procedure of sample preparation illustrated in the Supplemental Fig. S9A). A comparison of differential gene expression in *endu-2(−)* and *endu-2*(+) backgrounds suggested a similar numbers of transcripts as up- and down-regulated by *endu-2* (fold-change >2) (Fig. 5A and Supplemental Tab. S2). Among the 258 transcripts down-regulated in *endu-2(lf)* animals, germline expressed genes (n=206) were over-represented (Supplemental Table S2). qPCR quantifications of two selected genes *cav-1* and *trsc-1* confirmed our microarray result and revealed that their expression levels strongly decreased in *endu-2* mutants compared with wild type animals at 25°C (Supplemental Fig. S10C and S10D). However, neither a germline expressed *cav-1::GFP* reporter nor smFISH staining of *trcs-1* confirmed significantly altered germline expression *in situ* (Supplemental Fig. S10A, S10B and S10D). As the gonads of *endu-2(lf)* mutants at 25°C became significantly smaller (Supplemental Fig. S9B), we speculate that apparent down-regulation of germline mRNA within the transcriptome data set may be caused by reduced sample size of the gonad in *endu-2* mutants.

**Fig. 5.**
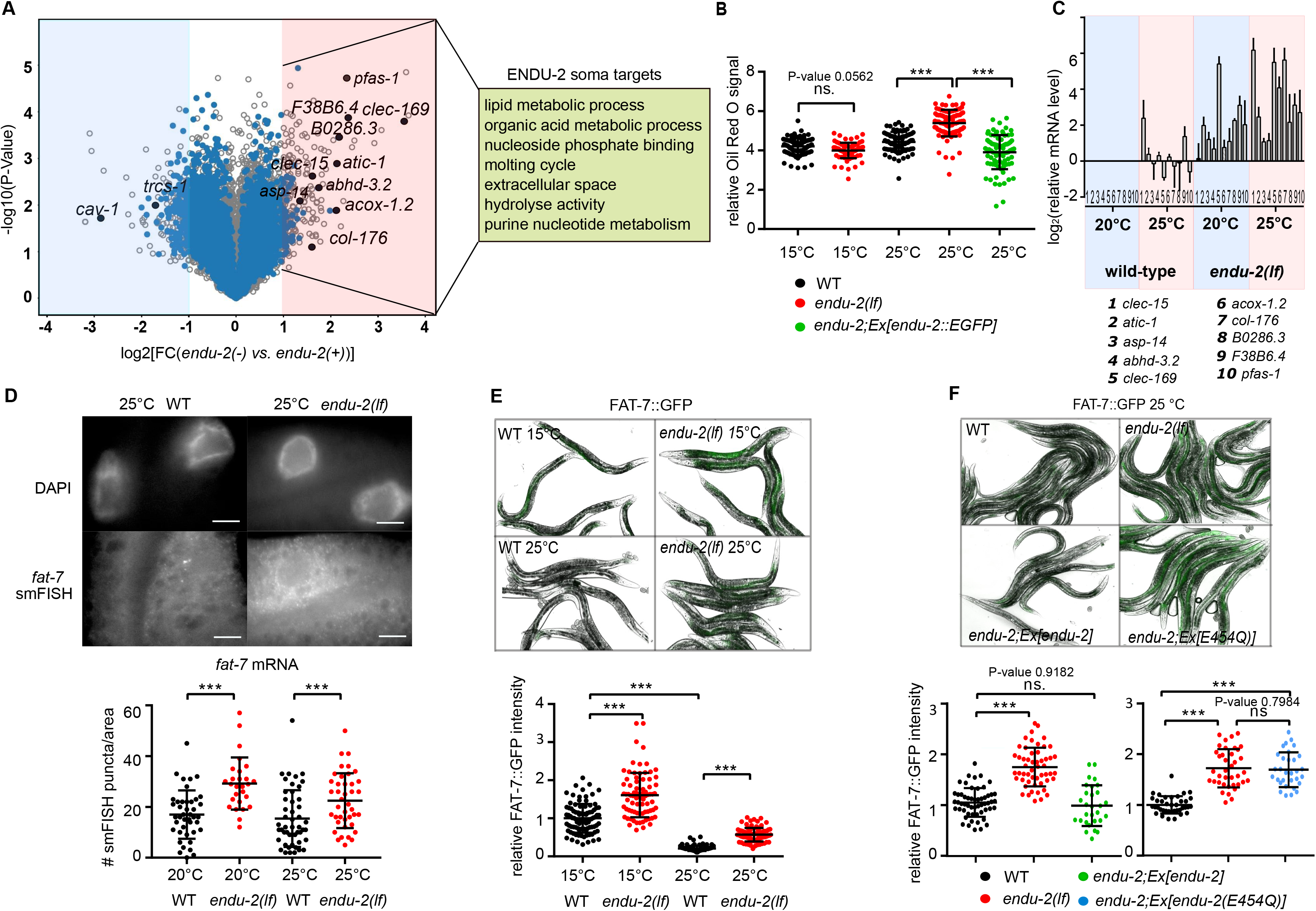
ENDU-2 negatively regulates mRNA abundance in the soma. **A.** Volcano blot of microarray results comparing *endu-2(tm4977)* mutant and *endu-2(tm4977)* animals carrying the rescuing *endu-2::EGFP* transgene at 25°C. N = 3 biological replicates. The blue dots mark transcripts bound by ENDU-2. Transcripts with blue and black label were ENDU-2 associated according to the RIP-Seq result and those with black labels were verified with q-PCR. **B.** ENDU-2 negatively regulates lipid content at 25°C. Shown are pooled data of quantification of relative Oil Red O stained signal in wild type, *endu-2(tm4977)* as well as *endu-2(tm4977);byEx1375[endu-2::EGFP]* day one adult animals at different temperatures. N=3 replicates. Statistical test with One-way ANOVA. **C.** qPCR to quantitate selected ENDU-2 target mRNAs in the soma at 20°C and 25°C. Data are mean ± SD, N= three biological replicates. **D.** Fluorescent images of smFISH staining of the ENDU-2 target *fat-7* mRNA and quantification of the smFISH stained mRNA foci per examined area in wild type and *endu-2(tm4977)* animals. Scale bar 10 μm. Data are mean ± SD of pooled data from three biological replicates. Statistical test with One-way ANOVA. **E.** Fluorescent images and quantification of fluorescence intensity of FAT-7::GFP in wild type and *endu-2(tm4977)* day one adult animals at different temperatures. Shown are pooled data from three biological replicates. Statistical test with One-way ANOVA. **F**. Fluorescent images and quantification of fluorescence intensity of FAT-7::GFP in wild type, *endu-2(tm4977)*, *endu-2(tm4977);Ex[endu-2(wt)]* and *endu-2(tm4977);Ex[endu-2(E454Q)]* day one adult animals at 25°C. Shown are pooled data from three biological replicates. Statistical test with One-way ANOVA. *** in this Figure means P value < 0.0001.

Therefore, we put our focus on genes that were up-regulated in *endu-2(lf)* back ground. 32% (76 out of 237) of these transcripts were candidates for direct ENDU-2 targets, since they had been co-immunoprecipitated with ENDU-2 (Supplemental Tab. S1). In addition, the vast majority of them (62 out of 76) did not have increased expression in the gonad of *endu-2(lf)* animals (see the next chapter, Supplemental Table S1). This suggests that, at minimum, these 62 transcripts are down-regulated by ENDU-2 in the somatic tissues. GO term analyses implicated these somatic ENDU-2 targets in regulation of various metabolic processes (Fig. 5A and Fig. S9C). Consistently, we found loss of *endu-2* resulted in increased lipid content at 25°C but not at 15°C (Fig. 5B). Taken together, these results indicate a putative regulatory role of ENDU-2 in limiting abundance of the mRNAs that are primarily involved in metabolic functions in the soma.

As some aspects of the *endu-2(lf)* phenotype were temperature dependent, we asked whether ENDU-2 regulates mRNA abundance in response to temperature alterations. To test this, we performed qPCR to quantitate ten selected mRNAs which were bound and down-regulated by ENDU-2. We found that mRNA levels of these targets were up-regulated in *endu-2* mutant background, compared to wild type, already at standard growth temperature (20°C) and were even more abundant at 25°C (Fig. 5C). Increased temperature, on the other hand, did not strongly affect the abundance of most of these mRNAs in wild type background. These results suggest a temperature dependent negative influence of ENDU-2 on the levels of these mRNA targets. We also performed smFISH to inspect the influence of ENDU-2 on the mRNA level of yet another target *fat-7*. *fat-7* mRNA was only detected in the intestine and *endu-2(−)* animals had higher *fat-7* transcript levels than wild type at both 20°C and 25°C (Fig. 5D). Moreover, we used a *fat-7::GFP* translational fusion reporter to monitor FAT-7::GFP protein level. Wild type animals showed reduced FAT-7::GFP expression at 25°C *vs.* 15°C (Fig. 5E). *endu-2(lf)* displayed stronger FAT-7::GFP expression at both temperatures. Furthermore, transgenic expression of *endu-2(wt)* but not the *endu-2(E454Q)* transgene (that has lost RNA cleavage activity) restored decreased FAT-7::GFP expression (Fig. 5F). We conclude that ENDU-2 mediated mRNA-cleavage is required for decreasing expression of at least some of its somatic target genes, such as *fat-7*.

### ENDU-2 prevents misexpression of soma specific genes in the germline

To investigate germline transcriptomes regulated by ENDU-2, we isolated gonads from wild type and *endu-2(lf)* animals which had been grown at 25°C from L1 stage, and sequenced two biological replicates of polyadenylated RNA from each strain. A gene with normalized reads of RPKM > 1 was scored as expressed. We identified 8,356 expressed genes in the gonad of wild type animals at 25°C, among which 88% (7,365) have been reported as expressed in the wild type gonad at 20°C (Ortiz et al., 2014). The following DESeq2 analysis showed that up-regulation of mRNA levels was the prevalent change in the gonad of *endu-2* mutants (422 up-regulated, 35 down-regulated, FC>2, P value<0.05; Fig. 6A, Supplemental Tab. S3). And only genes up-regulated in *endu-2(lf)* background were enriched in ENDU-2 binding, as 218 out of 422 up-regulated genes (P-value < 0.0001, Chi-Square test) and 11 out of 35 down-regulated genes (P-value = 0.7675, Chi-Square test) were co-immunoprecipitated with ENDU-2. We considered them as direct targets of ENDU-2 and grouped them in two classes, depending on whether they are up- (class I) or down-regulated (class II) in *endu-2(lf)* background (Fig. 5B). Expression of a class I target is repressed while class II genes are activated by wild type ENDU-2 activity. Consistently, we noticed that most of the class I, but not the class II target genes were not, or very lowly, expressed in the gonad of wild type animals (Fig. 5C) and tissue enrichment analysis suggested that the class I targets are enriched expressed in the somatic tissues (intestine, neurons, muscles, somatic gonad and pharynx). GO term enrichment analysis revealed that the major function of class I targets are involved in immune and defense responses (Fig. 5B). We wondered whether the class I target genes were preferentially soma specific genes that are misexpressed in the germline in the absence of ENDU-2. To validate this assumption, we combined our wild type gonad transcriptome with a previous published whole animal’s transcriptome at 25°C (Gomez-Orte et al., 2018) and calculated a soma enrichment factor (SEF) for each transcript to estimate the expression ratio between the soma and the gonad. Class I but not class II targets had on average significantly higher SEF value than the total mRNAs (Fig. 6D), suggesting that the class I genes are predominantly expressed in the soma but only expressed at low level in the germline in wild type animals. As maintenance of germline immortality by ENDU-2 does not require its RNA cleavage activity, we additionally sequenced mRNA from isolated gonads of *endu-2(lf)* mutant carrying *endu-2(E454Q)::EGFP* rescue transgene. Expressing *endu-2(E454Q)::EGFP* was sufficient to repress germline expression of most all the class I targets (Fig. 6E and 6F). In summary, our data indicate the inhibitory role of ENDU-2 to prevent germline expression of soma-specific genes is primarily independent of its RNA cleavage activity.

**Fig. 6.**
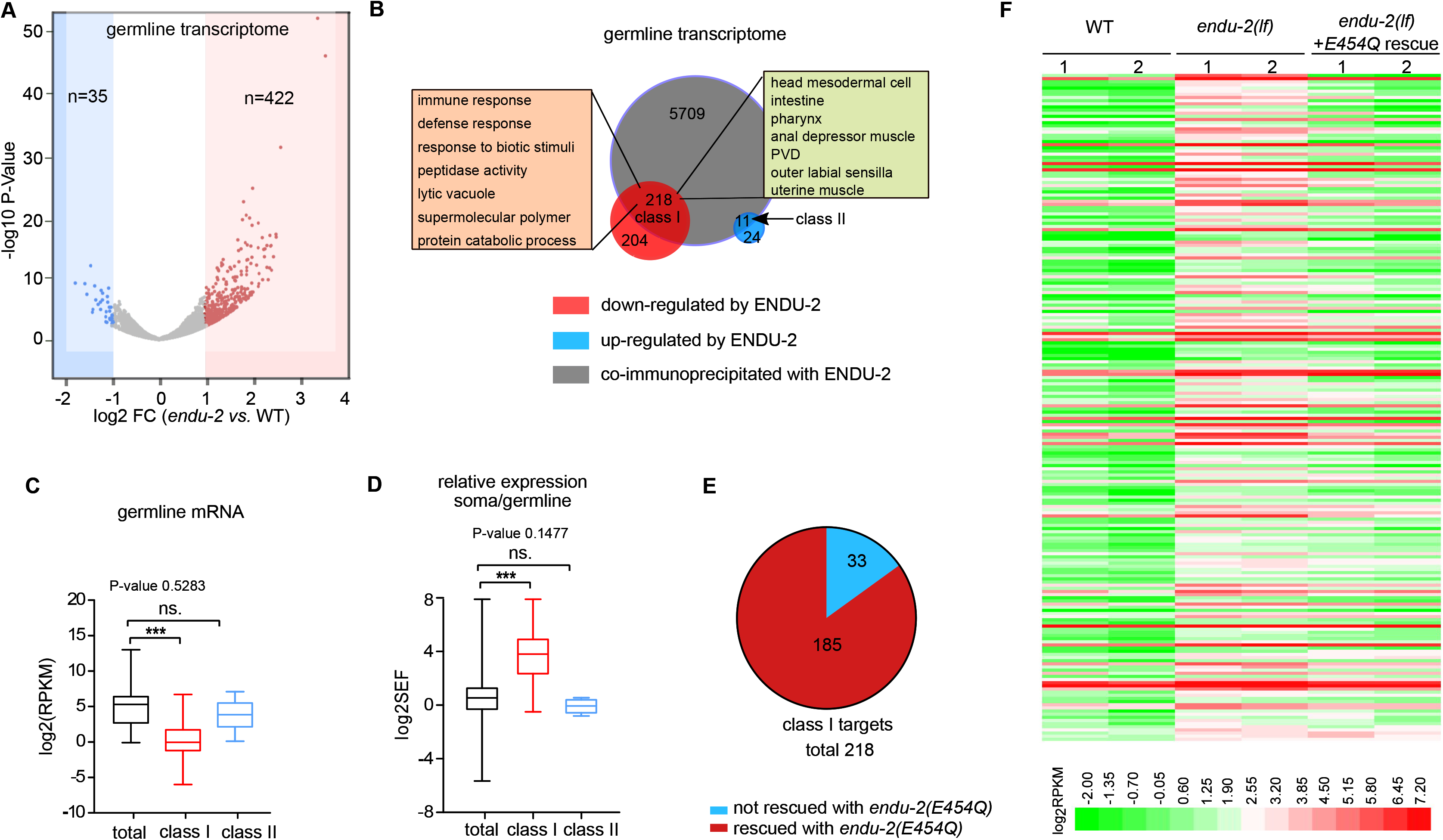
ENDU-2 prevents misexpression of soma specific genes in the germline. **A.** Volcano plot of gonad expressed genes in *endu-2(tm4977)* relative to wild type day one adult animals grown up at 25°C. 2 biological replicates for each strain. **B.** Comparison of the total transcripts co-immunoprecipitated with ENDU-2 and ENDU-2 regulated genes in the gonad. **C.** Comparison of expression levels (RPKM) of total gonad-expressed genes, ENDU-2 class I and class II targets in wild type animals. Shown are Box-Plot of Min/Max whiskers. Statistical test with One-way ANOVA. **D.** Comparison of relative expression between the soma and the gonad of the total transcripts, ENDU-2 class I and class II targets in wild type animals. Shown are Box-Plot of Min/Max whiskers. Statistical test with One-way ANOVA. **E.** The majority of the alerted expression of the germline class I targets is rescued by *endu-2(E454Q)::EGFP* transgene. **F.** Heat map of ENDU-2 repressed germline genes in wild type, *endu-2(tm4977)* and *endu-2;Ex[endu-2(E454Q)::EGFP]* animals. *** in this Figure means P value < 0.0001

## Discussion

The task of germline is to maintain pluripotency and immortality for an accurate transmission of genetic and epigenetic information between generations. Despite of emerging studies about molecular mechanisms protecting the germline from the impacts of stress, one obvious open question is whether the germline responds directly to environmental signals, such as nutrients or stress, or rather reacts to alterations in somatic signaling. A direct stress response seems to be less advantageous from an economical point of view, since this requires, in a pluripotent stem cell, the expression of an entire hierarchy of mechanisms, involving the sensation of stress, signaling, as well as responsive pathways to enable a dynamic and rapid response to a wide range of environmental cues. In contrast, the second alternative would more beneficial as differentiated somatic cells are specialized to sense environmental changes. In such a case, molecular messengers, such as secreted hormones and ligands, are required to shuttle between somatic and germline tissues to adjust a proper response in the germline. Previous studies have revealed that microRNA could act as such messengers mediating communication between soma and germline (Maciel and Mansuy, 2019; Sharma et al., 2018). Here we show secretion of a conserved endoribonuclease ENDU-2 from the soma prevents misexpression of soma-specific gene in the germline and preserve germline immortality at elevated temperature. This finding, together with another recent study reporting regulation of germline proliferation by intestinal ENDU-2 in response to thymidine imbalance (Jia et al., 2020), strongly suggests ENDU-2 as a crucial molecule mediating non-cell-autonomous stress responses.

Our data implicate ENDU-2 in enabling long range communication between cells via its secretion signal peptide (Fig. 2B). Whereas an earlier study had claimed neurons and muscles as the tissues primarily expressing *endu-2* (Ujisawa et al., 2018), our data, consistent with the report from Jia et al. (2020), suggest intestine as the most important organ to produce ENDU-2, whereas expression in neuronal, muscular and somatic gonad tissues is rather weak (Fig. 2A and Supplemental Fig. S3). We compared, for this analysis, both RNA and protein levels of ENDU-2 as well as different ENDU-2 variants with truncations in the N-terminal secretion signal peptide (Δ_ss_ENDU-2 and SS_sel-1_:: Δ_ss_ENDU-2). We could demonstrate that the tissues containing ENDU-2 protein differ from those that harbor *endu-2* mRNA and presence of a secretion signal peptide is necessary and sufficient to target ENDU-2 to the secretory pathway to reach distant cells.

Our data strongly suggests that ENDU-2 production and function may occur in different tissues, and that secretion of ENDU-2 allows the control of mRNA abundance in the distance. A simple assumption would be that ENDU-2 has similar functions in any of its target tissues, no matter from which cells it is expressed. This seems to be the case for some activities mediated by ENDU-2. For example, the egg-laying defect observed as a consequence of abnormal vulva development in an *endu-2(lf)* mutant is rescued by transgenic *endu-2* expression in the muscle, intestine or neuron (data not shown). In contrast, only neuronally or intestinally expressed ENDU-2 was able to rescue germline immortality (Fig. 3C), indicating existence of functional difference of ENDU-2 from distinct origins. Tissue-specific interactors or modifiers of ENDU-2 are probably crucial to specify its individual activities. It is currently not known whether there are isoform-specific functions of ENDU-2, or whether ENDU-2 protein is modulated by protein modifications.

We distinguish two activities of ENDU-2: mRNA-binding and mRNA-cleavage. RIP-Seq and transcriptomic results demonstrate that the levels of only about 10% of the mRNAs to which ENDU-2 binds change in *endu-2(lf)* mutants at elevated temperature (Fig. 4–6), suggesting that ENDU-2 is able to discriminate its regulated targets from the bound transcripts. Such scenarios have also been proposed for SMG-2/UPF1, the core factor of the non-sense mediated decay (NMD) complex that degrades only a small fraction of the RNAs it binds to (Muir et al., 2018). ENDU-2 represses expression of its somatic target genes probably via direct RNA degradation while it utilizes primarily RNA-binding but not cleavage activity to prevent misexpression of soma-specific genes in the germline, suggesting rather an indirect mechanism of ENDU-2 in the germline to restrict its target mRNA abundance. Why such different layers of regulatory mechanisms exist is currently unknown. We speculate that one reason might be association of ENDU-2 with distinct protein complexes in the soma and germline. Future study focusing on dissecting tissue specific mechanisms will help to elucidate how ENDU-2 coordinates gene expression in the soma and germline upon environmental stimuli.

*Xenopus* EndoU was suggested to both control snoRNA biogenesis in the nucleolus and cleave mRNAs in the cytosol (Gioia et al., 2005; Schwarz and Blower, 2014). Mouse EndoU has been reported to regulate peripheral B cell survival via reducing c-Myc mRNA level (Poe et al., 2014). Inactivation of neuronal Drosophila DendoU causes neurotoxicity partially by down-regulation of dTDP-43 (Laneve et al., 2017). SARS and SARS-Covid-2 endoribonucleases Nps15/NendoU have been implicated in virus replication as well as limitation of host innate immune response, although the detailed functions are still enigmatic (Kim et al., 2020). Our work here suggests that *C. elegans* ENDU-2, in response to environmental stimuli such as temperature elevation, can exert its function in tissues different from the ones in which its gene is expressed. Therefore, ENDU-2 is a novel candidate for transmitting environmental signals across tissue boundaries, most notably involving signaling to the germline. A secretion signal peptide is also present in the human EndoU homolog that is expressed in the placenta and detected in the serum (Inaba et al., 1980). Therefore, the potential conservation of its non-cell-autonomous functions could open a new research area to study intercellular communication. EndoU, thus, could be involved in a mechanism by which environmental influences and experiences of the somatic tissues are transmitted into the reproductive system.

## Methods

### Strains

The *C. elegans* N2 (Bristol) strain was used as wild type in all experiments in this study. Mutant strains: KHR83 *endu-2(tm4977),* outcrossed 8x with our N2. BR7130 *endu-2(by188)*, NL3511 *ppw-1(pk1425),* TX20 *oma-1(zu405)*, BR7510 *oma-1(zu405);endu-2(tm4977)*, BR8649 *oma-1(zu405);hrde-1(tm1200),* BR7205 *endu-2(by190[endu-2::EGFP])*, BR7089 *byEx1315[endu-2P::EGFP;rol-6(su1006)]*, BR8657 *endu-2(tm4977);byEx1814[endu-2P::endu-2::EGFP::_endu-2_3’UTR;rol-6(su1006);]*, BR7295 *endu-2(tm4977);byEx1375[endu-2P::endu-2::EGFP*;*myo-2P::mCherry*], BR5402 *byEx749[unc-119P::EGFP;rol-6(su1006)]*, BR8672 *byEx1821[unc-119P::SS_endu-2_::EGFP;rol-6(su1006)]*, BR7827 *endu-2(tm4977)*;*byEx1551[vha-6P::endu-2::EGFP::3xFlag;myo-2P::mCherry]*, BR7332 *endu-2(tm4977);byEx1379[fos-1P::endu-2::EGFP;myo-2P::mCherry]*, BR8551 *endu-2(tm4977);byEx1795[unc-119P::endu-2::EGFP::3xFlag;rol-6(su1006)]*, BR8662 *endu-2(tm4977);byEx1816[myo-3P::endu-2::EGFP::3xFlag;myo-2P::mCherry],* BR7512 *endu-2(tm4977);byEx1449[endu-2P::Δ_ss_endu-2::EGFP;rol-6(su1006);myo-2P::mCherry]*, BR8747 *endu-2(tm4977);byEx1847[endu-2P::Δ_ss_endu-2::EGFP::_endu-2_3’UTR;myo-2P::mCherry]*, BR8821 *endu-2(tm4977);byEx1875[endu-2P::SS_sel-1_::Δ_ss_endu-2::EGFP; myo-2P::mCherry],* BR7680 *endu-2(tm4977);byEx1492[endu-2P::endu-2(E454Q)::EGFP::3xFlag;myo-2P::mCherry]*, BR7683 *endu-2(tm4977);byEx1495[endu-2P::endu-2(E460Q)::EGFP::3xFLAG;myo-2P::mCherry]*, DMS303 *nIs590[fat-7p::fat-7::GFP*+*lin15*(+)], BR8317 *endu-2(tm4977);nIs590[fat-7p::fat-7::GFP*+*lin-15*(+)], BR8754 *endu-2(tm4977);nIs590[fat-7p::fat-7::GFP*+*lin-15(*+)*];byEx1853[endu-2P::endu-2(E454Q);myo-2P::mCherry*], BR8775 *endu-2(tm4977);nIs590[fat-7p::fat-7::GFP+lin-15*(+)*];byEx1860[endu-2P::endu-2;myo-2P::mCherry]*, RT688 *unc-119(ed3);pwIs28[pie-1p::cav-1::GFP(7)*+*unc-119(*+)], BR8358 *endu-2(tm4977);pwIs28[pie-1p::cav-1::GFP(7)*+*unc-119(*+)]. For RIP-Seq the following strains were used: BR7802 *endu-2(tm4977);byIs240[endu-2P::endu-2(E460Q)::EGFP::3xFLAG;myo-2p::mCherry]*, BR7803 *endu-2(tm4977);byIs241[endu-2P::endu-2(E454Q)::EGFP::3xFLAG;myo-2P::mCherry]*, BR7205 *endu-2(by190[endu-2::EGFP])* and *Ex[ife-2P::GFP]* (as IP control, gift from Tavernarakis Lab). For the RNA cleavage assay, BR8311 *endu-2(tm4977);byIs267[endu-2P::endu-2::EGFP;myo-2::mCherry]* and BR7803 *endu-2(tm4977);byIs241[endu-2P::endu-2(E454Q)::EGFP::3xFlag;myo-2P::mCherry]* were used in additional to N2 and *endu-2(tm4977)* strains. *endu-2(tm4977)* allele was used for all experiments, if not noted otherwise. Except for the Fig. 1A and Supplemental Fig. S2, all the *endu-2(lf)* mutants were the granddaughter generation (G2) from a *endu-2(tm4977)* carrying transgenic *endu-2* rescue strains whose parents (G1) had lost the rescue transgene. Information for extrachromosomal transgenic strains generated in this study is summarized in the Supplemental Tab. S5.

### Plasmids

To construct an EGFP transcriptional fusion reporter of *endu-2* (pBY3798), a 4691bp genomic fragment upstream of the *endu-2* ATG was PCR amplified and inserted into pEGFP-N1 with *Eco47III/BglII* sites. For translational fusion reporters, 5487 bp *endu-2* genomic region was cloned into pBY3798 with *BglII/SmaI* sites to receive *endu-2P::endu-2::EGFP (*pBY3800), the *endu-2* 3’UTR was inserted into pBY3800 at *NotI* site to receive *endu-2P::endu-2::EGFP:: _endu-2_3’UTR* (pBY4137). *Δ_ss_endu-2::EGFP* (pBY3843) and *Δ_ss_endu-2::EGFP::_endu-2_3’UTR* construct (pBY4172) was generated by removing sequences encoding the N-terminal 2-19 amino acids from pBY3800 and pBY4137, respectively. The *endu-2P::SS_sel-1_::Δ_ss_endu-2::EGFP* construct (pBY4194) was generated by insert the sequence encoding the 1-20 amino acids of *sel-1* to the pBY3843 with *BglII* site. A *endu-2P::3xFlag::endu-2::EGFP*:: *_endu-2_3’UTR* expressing construct (pBY4138) was generated by inserting 3xFlag encoding sequence and *endu-2* genomic region into pBY3798 with *BglII* and *BglII/SmaI* sites, respectively. The 3’UTR of *endu-2* was inserted via Gibson ligation. The *unc-119P::EGFP* expressing construct (pBY2941) was made by fusing the 2.1 kb *unc-119* promoter region into pEGFP-N1 with *Eco47III/BglII* sites. The *unc-119P::SS_endu-2_::EGFP* construct pBY4148 was generated by inserting the first 57 nucleotides of *endu-2* cDNA into pBY2941 with *BglII/AgeI* sites. For expressing recombinant ENDU-2::EGFP in HEK 293T cells, wild type *endu-2* cDNA was cloned into a 3xFlag containing pEGFP-N1 with *XhoI/SmaI* (pBY3878). Site-directed mutagenesis was performed to receive *endu-2(E454Q)::EGFP::3xFlag* (pBY3894) and *endu-2(E460Q)::EGFP::3xFlag* mutants (pBY3895). The fragments *endu-2::EGFP::3xFlag from pBY3878, endu-2(E454Q)::EGFP::3xFlag* from pBY3894 and *endu-2(E460Q)::EGFP::3xFlag* from pBY3895 were inserted into pBY3798 to receive constructs for generating *endu-2P::endu-2::EGFP::3xFlag* (pBY3892), *endu-2P::endu-2(E454Q)::EGFP::3xFlag* (pBY3897) and *endu-2P*::*endu-2(E460Q)::EGFP::3xFlag* (pBY3898) reporter strains. *EGFP::3xFlag* fragments in pBY3892 and pBY3897 were removed to receive pBY4188 *endu-2P*::*endu-2* and pBY4066 *endu-2P*::*endu-2(E454Q)*. Somatic gonad specific expression was achieved via PCR amplifying and insertion of *endu-2* cDNA into a *fos-1P::GFP* plasmid (a gift from David Sherwood lab) with *SalI* and *SmaI* sites (pBY3833). For intestinal specific expression, a 1.2 kb *vha-6* promoter was inserted at *Eco47III/XhoI* sites into pBY3878 to receive *vha-6P::endu-2::EGFP::3xFlag* (pBY3937). A 2.1 kb *unc-119* promoter was inserted into pBY3878 at *Eco47III/BglII* sites to receive *unc-119P::endu-2(cDNA)::EGFP::3xFlag* (pBY4127) for neuronal expression. A 2.4 kb *myo-3* promoter region was cloned into pBY3878 to receive *myo-3P::endu-2::EGFP::3xFlag* (pBY4135) for muscular specific *endu-2* expression.

### Scoring of Mrt phenotype

Mrt phenotype was examined by quantification of brood size by separating 15 L4 animals on single plates and counting number of progeny per animal, or transferring six L4 larvae onto new agar plates and scoring percentage of the fertile animals 36 hours after the L4 stage in the next generation. Both *endu-2(tm4977)* and *endu-2*(*by188)* were further outcrossed 4x with N2 before scoring Mrt phenotype for Fig. 1A and Supplemental Fig. S2A. For brood size recovery at 15°C, wild type and outcrossed *endu-2(tm1977)* animals were maintained at 25°C for 6 generations until *endu-2(tm4977)* showed strong sterile phenotype. 15 L1-L2 animals from the 6th generation were shifted to 15°C and designated as G1. For Mrt rescue experiment, *endu-2(tm4977)* progenies of *endu-2(tm4977);byEx1375[endu-2::EGFP;myo-2P::mCherry], endu-2(tm4977);byEx1449[endu-2P::Δ_ss_endu-2::EGFP;myo-2P::mCherry], endu-2(tm4977);byEx1875[endu-2P::SS_sel-1_::Δ_ss_endu-2::EGFP;myo-2P::mCherry], endu-2(tm4977);byEx1488[endu-2P::endu-2(E454Q)::EGFP::3xFLAG;myo-2P::mCherry]* and *endu-2(tm4977);byEx1488[endu-2P::endu-2(E460Q)::EGFP::3xFLAG;myo-2P::mCherry]* animals (P0) were isolated and designated as G1.

### Germline proliferation

Number of germ nuclei in the proliferation zone was quantified as described in a previous study (Qi et al., 2017).

### Lifespan

Lifespan at 20°C was initiated at L4 stage. As *endu-2(tm4977)* animals display strong egg-laying defect due to abnormal vulval development, agar plates containing 200μM FUDR were used to during the first seven days of adulthood to avoid internal hatching.

### Antibody staining

Antibody staining was performed with day one adult animals as described (Crittenden and Kimble, 2009). A monoclonal mouse anti-GFP antibody (Roche, Nr. 11814460001) was used to detect ENDU-2::EGFP in the gonad.

### *oma-1* RNAi inheritance assay

*oma-1* RNAi was initiated at L1 stage at 20°C (restrictive temperature) and these animals were designated as P0. Six P0 day one adults animals were transferred onto OP50 seeded agar plates and incubated at 20°C for scoring the embryonic lethality over generations. Embryonic lethality was examined by transferring of about 100 embryos onto an OP50 seeded agar plate and quantifying the number of hatched animals after 48 hours.

### Oil Red O staining to quantify content of body fat

Worms were fixed in 500μl 60% isopropanol. After removal of supernatant, 500μl freshly filtered ORO working solution was added to stain worms at 25°C in a wet chamber overnight. Animals were washed once with 1ml 0.01% Triton-x100 containing M9 buffer and responded in 250μl M9-Triton-x100 buffer. Stained worms can be stored at 4°C for at least one month. For quantification, images were recorded with color camera. The original images were spitted into red and blue channels and the level of Oil-Red-O was quantified by determining the excess intensity in the red channel in comparison to the blue channel. The mean fatness per worm was calculated as the total intensity within stained regions normalized by the area of the worm regions. Oil Red O working solution was prepared as follows: 0.5 g of Oil Red O powder was dissolved in 100 ml isopropanol solution and equilibrated for several days. The solution was then freshly diluted with 40% water for a 60% stock and allowed to sit overnight at room temperature and filtered using 0.2 mm filters.

### RIP-Seq

To isolate ENDU-2 associated RNAs, animals of mixed stage were lysed in Lysis-NP-40 buffer (150 mM NaCl,50 mM Tris pH8.0, 1% NP40 + Protease Inhibitor+100 μl/ml RNase Inhibitor) with SilentCrusher (x30 sec. 75000 rpm). Lysates were clarified by centrifuging at 14k rpm for 15 min. Supernatants were pre-cleared with Dynal Protein A magnetic beads (Invitrogen) and incubated with anti-GFP antibody (Abcam) for 1 hour at 4°C, followed by incubation with Dynal Protein A Magnetic Beads for 2 h at 4°C. Beads were washed 3 times with Lysis-NP-40 Puffer before DNase I digestion (100 μl 1x DNase buffer, 1 μl DNase I, 1 μl RNase inhibitor) at 37 °C for 30 minutes. 10 μl 50 mM EDTA was added to stop DNase digestion. RNA was extracted with 5 volumes Trizol reagent (Invitrogen), followed by isopropanol precipitation. Precipitates were resolved in RNase-free water for visualization on 12.5% denaturing acrylamide gel with SYBR Gold (Thermo Scientific). Random hexamer primers were used for preparing TruSeq3 reverse-stranded cDNA libraries for Illumina single-end 50bp sequencing. RIP-Seq to compare RNA binding efficiency of ENDU-2(wt), ENDU-2(E454Q) and ENDU-2(E460Q) at 15 °C and 25 °C were performed without cross-linking. RIP-Seq to determine RNA targets of ENDU-2 was carried out with ENDU-2(E454Q)::EGFP expressing animals raised at 15 °C with UV cross-linking.

### Analysis of RIP-Seq data

The sequencing data were uploaded to the European Galaxy Server at https://usegalaxy.eu, and all read quality controls, trimming, mapping and counting were performed through Galaxy (Afgan et al., 2018). Specifically, after trimming with Trimmomatic 0.36 (Bolger et al., 2014), we mapped the reads to the *C. elegans* genome using the Wormbase WS260 genomic sequence (ftp://ftp.wormbase.org/pub/wormbase/releases/WS260/species/c_elegans/PRJNA13758/c_elegans.PRJNA13758.WS260.genomic.fa.gz) and the WS260 canonical gene set (ftp://ftp.wormbase.org/pub/wormbase/releases/WS260/species/c_elegans/PRJNA13758/c_elega ns.PRJNA13758.WS260.canonical_geneset.gtf.gz) with the RNA aligner STAR 2.6.0b (Dobin et al., 2013) with default settings, then counted reads mapping to individual genes using feature Counts 1.6.2 (Liao et al., 2014). For a read to be counted as mapping to a particular gene, we required a minimum read mapping quality of 12 (-Q 12 option of feature Counts) and an overlap of at least 1 base between the read and any of the exons of the gene (--minOverlap 1).

The IP enrichment of a transcript was expressed as: FC=(IP RPM + 1)/(Mock IP RPM + 1), where RPM= number of reads mapped to a gene/(number of all mapped reads) × 10^6. The addition of 1 in the FC formula prevents infinite enrichment. ENDU-2 associated RNAs were defined as transcripts with both FC>=4 and RPKM >=1 in ENDU-2(E454Q) IP sample. RPKM is calculated as follows: RPKM = RPM/length of transcripts × 10^3.

### RNA-seq and data processing of isolated gonads

Animals were raised at 25°C from the L1 larval stage and the complete gonads were isolated 24 h after mid-L4 stage. Total RNA of the gonads was extracted with RNeasy® Mini Kit (Qiagen, Venlo, The Netherlands). Purification of poly-A containing RNA molecules, RNA fragmentation, strand-specific random primed cDNA library preparation and Single-read sequencing (50 bp) on an Illumina HiSeq 4000 were carried out by Eurofins Genomics. The RNA-seq results from two independent biological replicate of dissected gonads were uploaded to the European Galaxy Server at https://usegalaxy.eu. All read quality controls, trimming, mapping and counting were performed through Galaxy (Afgan et al., 2018) using protocols like described for RIP-Seq (above). The DESeq2 (Galaxy Version 2.11.40.6+galaxy1) was used to determine differentially expressed features from count tables of differential transcript abundances. The RPKMs throughout this study were normalized separately. Only the longest isoform of each gene was used as length of a transcript. To estimate genes with enriched expression in the soma, RNA-seq data from wild type animals raised at 25°C from a previous study (Gomez-Orte et al., 2018) were compared with our transcriptome from isolated gonads of wild type animals. The soma enrichment factor (SEF) was calculated with the formal SEF=(RPKM_whole animal_ +2)/(RPKM_gonad_ +2)]. Addition of factor 2 to the formal minimize large SEF value caused by low RPKM values.

### In *vitro.* RNA binding assay of recombinant ENDU-2 protein

The RNA binding assay was performed as described (Panda et al., 2016). *fat-7* and *trcs-1* mRNA were transcribed from complementary DNA (cDNA) with biotinylated uracil. The presence and purity of RNA were examined in an RNA gel. This biotin-labeled RNA was incubated with recombinant ENDU-2(wt)::EGFP::3xFlag, ENDU-2(E454Q)::EGFP::3xFlag and ENDU-2(E460Q)::EGFP::3xFlag proteins expressed in HEK 293T cells. The labeled RNA was captured and isolated using streptavidin beads. Proteins binding the RNA were then visualized via Western Blot.

### RNA cleavage assay

Pellets of animals raised at 15°C in mixed stages were washed twice with 1 ml 10 mM EGTA containing M9 buffer before resuspension in 0.7 ml NP-40 buffer. 3x Protease inhibitor was added to the buffer and worms were lysed with SilentCrusher (2×30”, 75,000 rpm). 5μl RNAsin Plus RNAse Inhibitor (40U/μl) was added to the worm lysate and centrifuged by 12,000 rpm for 15 min at 4°C. For the RNA cleavage assay 95 μl supernatant of the worm lysate with 200 μg total protein was incubated with 5 mM Ca^2+^, Mg^2+^, Mn^2+^ or 10 mM EGTA for 30 minutes at 7°C before total RNA was extracted with Trizol. 0.2 μg RNA was loaded on 10% 8M Urea acrylamide gel with SYBR Gold for visualization of RNA.

### Microarray

Both *endu-2(tm4977)* and *endu-2(tm4977);byEx1375[endu-2P::endu-2::EGFP;myo-2::mCherry]* animals used for microarray were granddaughter decedents of one single *endu-2(tm4977);byEx1375[endu-2P::endu-2::EGFP;myo-2::mCherry]* animals (stain preparation see Supplemental Fig. S8). Animals were raised at 25°C for 48 h after hatching and total RNA was purified by using RNeasy® Mini Kit (Qiagen, Venlo, The Netherlands). Microarray and the following data analysis were performed as described before (Qi et al., 2017; Ritchie et al., 2015).

### GO term enrichment analysis

GO term and tissue enrichment analysis was carried out with the online enrichment analysis tool (https://wormbase.org/tools/enrichment/tea/tea.cgi)(Angeles-Albores et al., 2016).

### Quantification of mRNA level via quantitative RT-PCR (q-PCR)

Total RNA was prepared from day one adult animals raised at 25°C or 20°C by using RNeasy® Mini Kit (Qiagen, Venlo, The Netherlands). *endu-2(tm4977)* animals were granddaughter decedents of *endu-2(tm4977);byEx1375[endu-2P::endu-2::EGFP;myo-2::mCherry]* animals. *act-4* was used as internal control. q-PCR primer sequences: *cav-1* forward aagtgctggtggagtagatgc, *cav-1* reverse tccgatagcgatgttctcttc, *clec-169* forward tggacacttgtaactgtgcaaga, *clec-169* reverse ttttcatttgaccactttagatcg, *F38B6.4* forward cccttgtcctcggaaattaga, *F38B6.4* reverse caggcccgaagatggtaat, *pfas-1* forward caagattggaagttccgaaga, *pfas-1* reverse ctcaatatccggacagtcgtc, *B0286.3* forward gacccgaaaacttggagctt, *B0286.3* reverse Tggaacgaatgagcacataatc, *atic-1* forward tgctacaaaaatgccagcag, *atic-1* reverse cgtttaaaggaagaccaacagc, *acox-1.2* forward agcggtgatctatggaagtga, *acox-1.2* reverse agctcagggatcttggacac, *abdh-3.2* forward atgacaccaccccaattgtc, *abdh-3.2* reverse tgctgtcatgagtacttcctgtg, *col-176* forward ccacaacctttgctccaatc, *col-176* reverse gagcacacttgatgcagtcg, *clec-15* forward tgcgccagaagggtattact, *clec-15* reverse cgcagaacgatctaaccaga, *asp-14* forward gctgcagttaccaacattacca, *asp-14* reverse agcaacgaagaaggttgagg.

### Single Molecule Fluorescent in Situ Hybridization (smFISH) and data analysis

38-48 probes of 20nt in length targeting each mature mRNA were designed with Stellaris RNA FISH probe designer on gene target specificity. Probes were produced, conjugated to CAL Fluor Red 610 and purified by Biocat (sequences of the probe sets are available in the supplemental Table S4). Wild type and *endu-2(tm4977)* L4 animals were selected and dissected 24 hours later for smFISH staining using a published protocol procedure (Lee et al., 2016). Images were acquired with an Image Z1 fluorescence microscope. Exposure times and acquisition settings were identical between replicates. mRNA puncta were quantified by using ImageJ 1.51s cell counter plugin. Regions of interest for acquisition were defined by nuclei DAPI staining.

### Statistical analysis

Experiments shown in this study were performed independently twice to four times. Details of the particular statistical analyses used, precise *P* values, statistical significance, number of biological replicas and sample sizes for all of the graphs are indicated in the figures or figure legends. *n* represents the number of animals tested, unless mentioned otherwise. N means the number of biological replicates. *** represents P-value < 0.0001.

## Data availability

Microarray data were submitted to the ArrayExpresss under accession number E-MTAB-7993. RIP-seq data were submitted to BioProject under BioProject ID PRJNA544611. RNA-seq data of isolated gonad were submitted to NCBI Sequence Read Archive with ID SUB7811674. The remaining data for RIP-Seq and RNA-seq of isolated gonad are located in Supplemental Table S1-S3. Correspondence and requests for materials should be addressed to B.R (baumeister@celegans) or Q. W.

## Acknowledgements

We thank N. Tavernarakis and the *Caenorhabditis elegans* Genetics Center which is funded by the NIH Office of Research Infrastructure Programs (P40 OD010440) for providing some strains; Dr. S Mitani for providing *endu-2(tm4977)* strain, Cornelia Habacher for sharing the RIP-Seq protocol, D. Sherwood for providing the *fos-1* promoter. We thank the staff of the Life Imaging Center (LIC) of the Albert-Ludwigs-University Freiburg for their microscopy resources. This work was funded by grants from the German Research Foundation (DFG) (SFB850, SFB1381), Germany’s Excellence strategy (CIBSS-EXC-2189 - Project ID 8 390939984) and the German Excellence Initiative (BIOSS - EXC294) to R.B.

## Author Contributions

W.Q. and R.B. designed the experiments and analyzed data. W.Q., E.D.v.G., F. X., Q. Z., L.L. and W.Y. performed the experiments. D.P. performed and analyzed microarray. W.Q. and W.M. analyzed RNA-Seq data. W.Q. and R.B. wrote the manuscript.

## Competing interest

The authors declare no competing financial interests.

Correspondence and requests for materials should be addressed to W.Q.

## Notes

### Competing Interest Statement

The authors have declared no competing interest.

