## Supplemental Data for "A secreted endoribonuclease ENDU-2 from the soma protects germline immortality in *C. elegans*"

**Fig. S1**

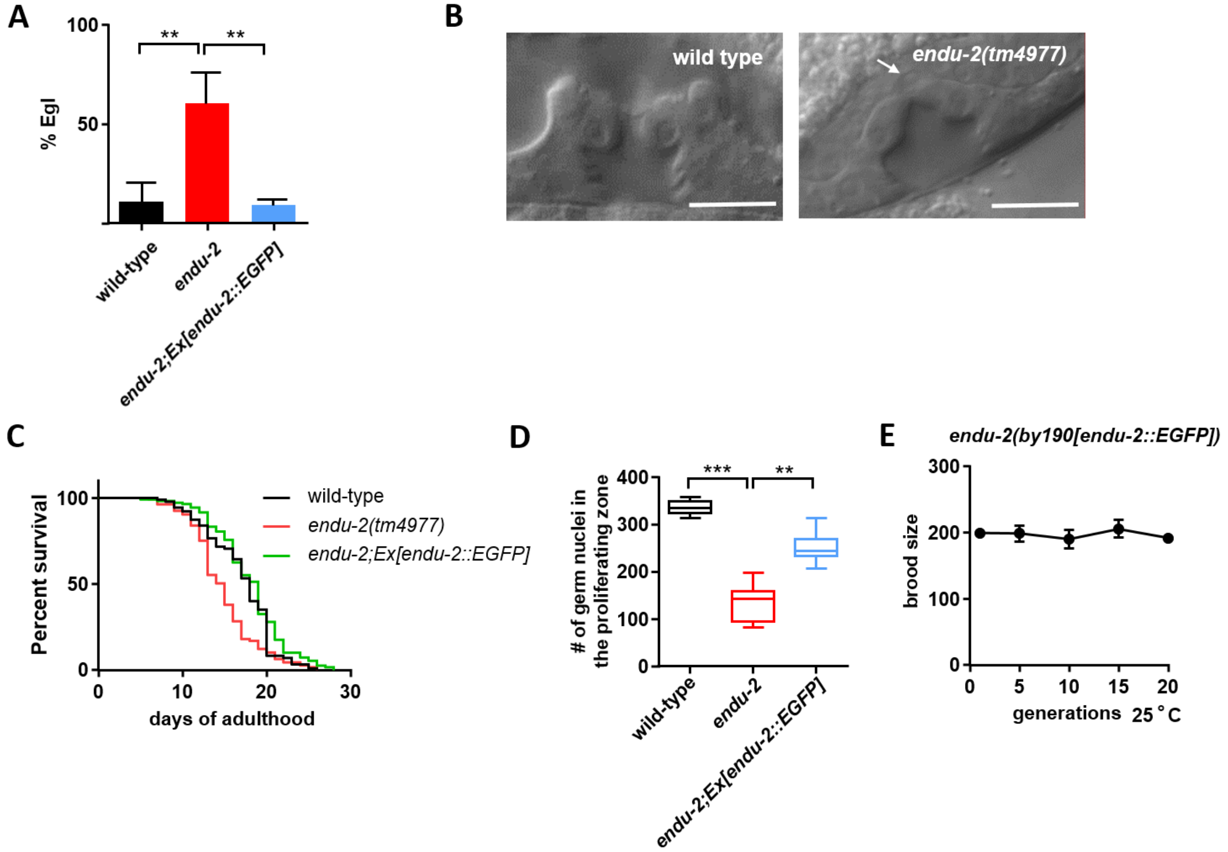


**Fig. S1 *endu-2* mutants phenotypes and the rescue with extrachromosomal *endu-2::EGFP* transgene.**

A) Extrachromosomal *endu-2::EGFP* transgene rescues egg-laying defects (Egl) of *endu-2(lf)* mutants at 20°C.

B) *endu-2(tm4977)* L4 animals show defective vulva development. The arrow points to the unfused anchor cell.

C) Extrachromosomal *endu-2::EGFP* transgene rescues short lifespan of *endu-2(lf)* mutants at 20°C. wild type: n=83, median survival 18 days, *endu-2(tm4977)*: n=105, median survival 15 days, *endu-2;Ex[endu-2::EGFP]*: n=107, median survival 19 days.

D) Extrachromosomal *endu-2::EGFP* transgene rescues reduced germline proliferation of *endu-2(lf)* mutants at 20°C.

E) The CRISPR EGFP knock-in animals do not show Mrt phenotype at 25°C.

All the rescue experiments in this figure were performed with BR7295 *endu-2(tm4977)X;byEx1375[endu-2P::endu-2::EGFP; myo-2P::mCherry]*.

This Figure is related to the main Figures 1-2.

**Fig. S2**

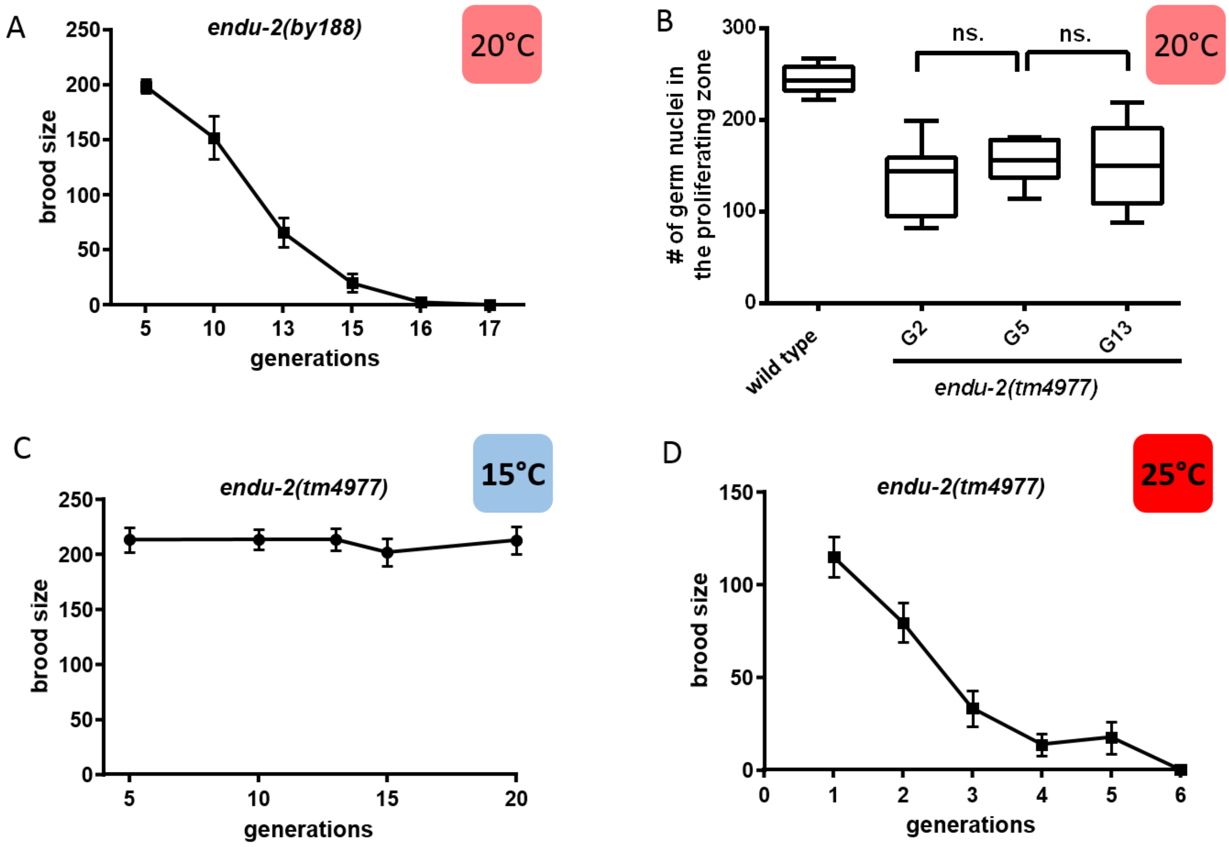


**Fig. S2 *endu-2(lf)* mutants have a temperature-sensitive Mrt phenotype.**

A) *endu-2(by188)* shows gradually reduced brood sizes over generations at 20°C.

B) Germline proliferation of *endu-2(tm4977)* mutants does not change over generations.

C) *endu-2(tm4977)* does not exhibit Mrt phenotype at 15°C.

D) Mrt phenotype of *endu-2(tm4977)* is enhanced at 25°C.

This Figure is related to the main Figure 1.

### Fig. S3

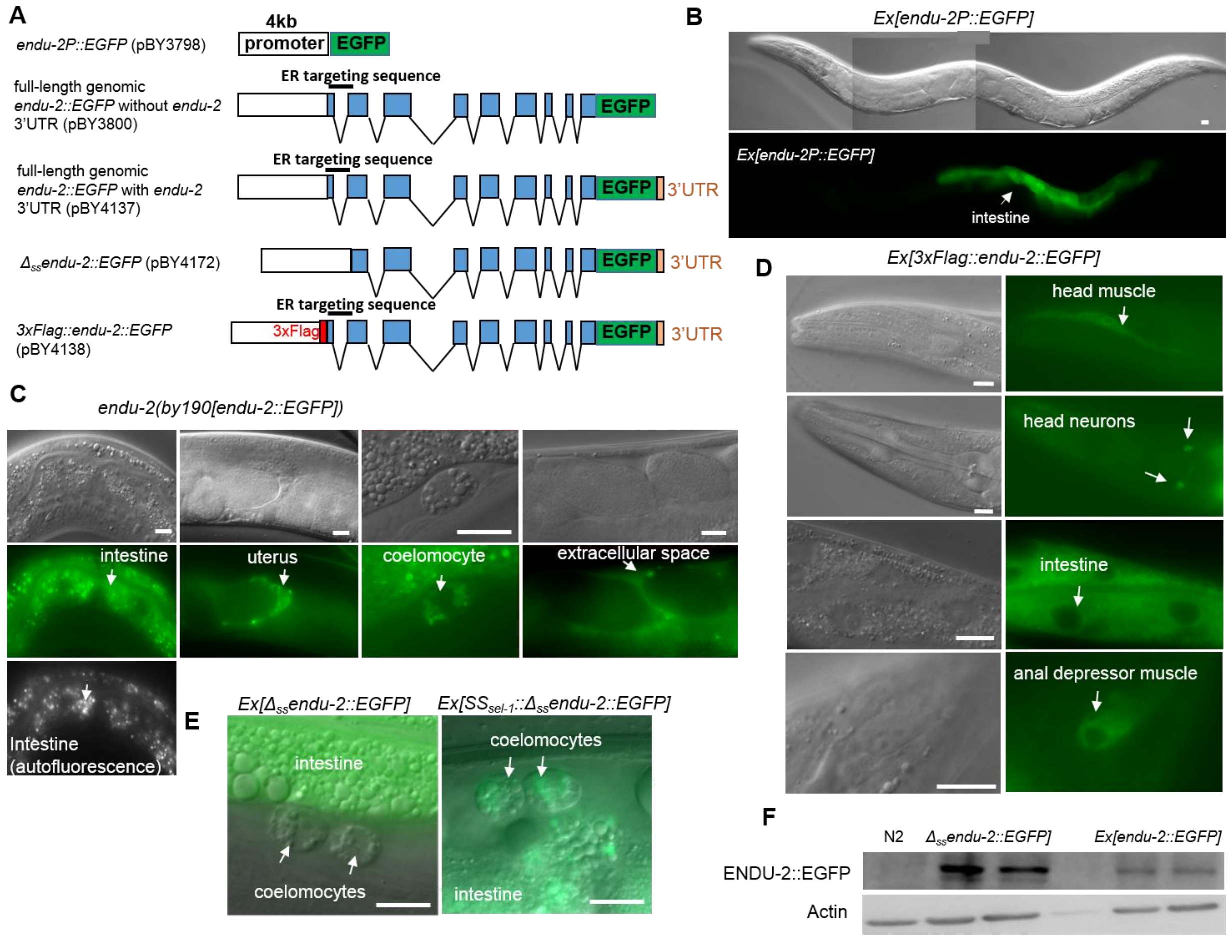


### Fig. S3 Expression pattern of *endu-2*

A) Summary of the transcriptional and translational fusion reporters of *endu-2* used for study expression pattern. All the transgenic strains generated for this study are listed in the Supplemental Tab. S5.

B) The EGFP transcriptional fusion reporter *byEx1315[endu-2P::EGFP]* shows intestinal expression.

C) The EGFP CRIPSPR-Cas9 knock-in strain *endu-2(by190[endu-2::EGFP])* displays weak ENDU-2::EGFP expression in the intestine, somatic gonad, coelomocyte and extracellular space.

D) Expression of 3xFlag::ENDU-2::EGFP (*byEx1805[endu-2P::3xFlag::endu-2::EGFP::*$_{endu\text{-}2}$*3'UTR]* in the intestine, head neuron, muscle cells in the head region and anal depressor muscle.

E) Intestinal expressed $\Delta_{ss}$ENDU-2::EGFP (BR7512 *endu-2(tm4977);byEx1449[endu-2P::*$\Delta_{ss}$*endu-2::EGFP]*) is not detected in the coelomocyte while fusion of the predicted secretion signal peptide (1-20 amino acids) of SEL-1 to the N-terminal of $\Delta_{ss}$ENDU-2::EGFP (BR8821 *endu-2(tm4977);byEx1875[endu-2P::*$SS_{sel\text{-}1}$*::*$\Delta_{ss}$*endu-2::EGFP]*) results in its localization in the coelomocyte.

F) Western Blot to detect transgenic expressed ENDU-2(wt)::EGFP and $\Delta_{ss}$ENDU-2::EGFP. Shown are two replicates with 10 day one adult animals of BR8657 *endu-2(tm4977);byEx1814[endu-2P::endu-2::EGFP::*$_{endu\text{-}2}$*3'UTR]* and BR8747 *endu-2(tm4977);byEx1847[endu-2P::*$\Delta_{ss}$*endu-2::EGFP::*$_{endu\text{-}2}$*3'UTR]* at 25°C. Wild type animals were used as negative control.

Scale bar 10 µm. This Figure is related to the main Figure 2.

**Fig. S4**

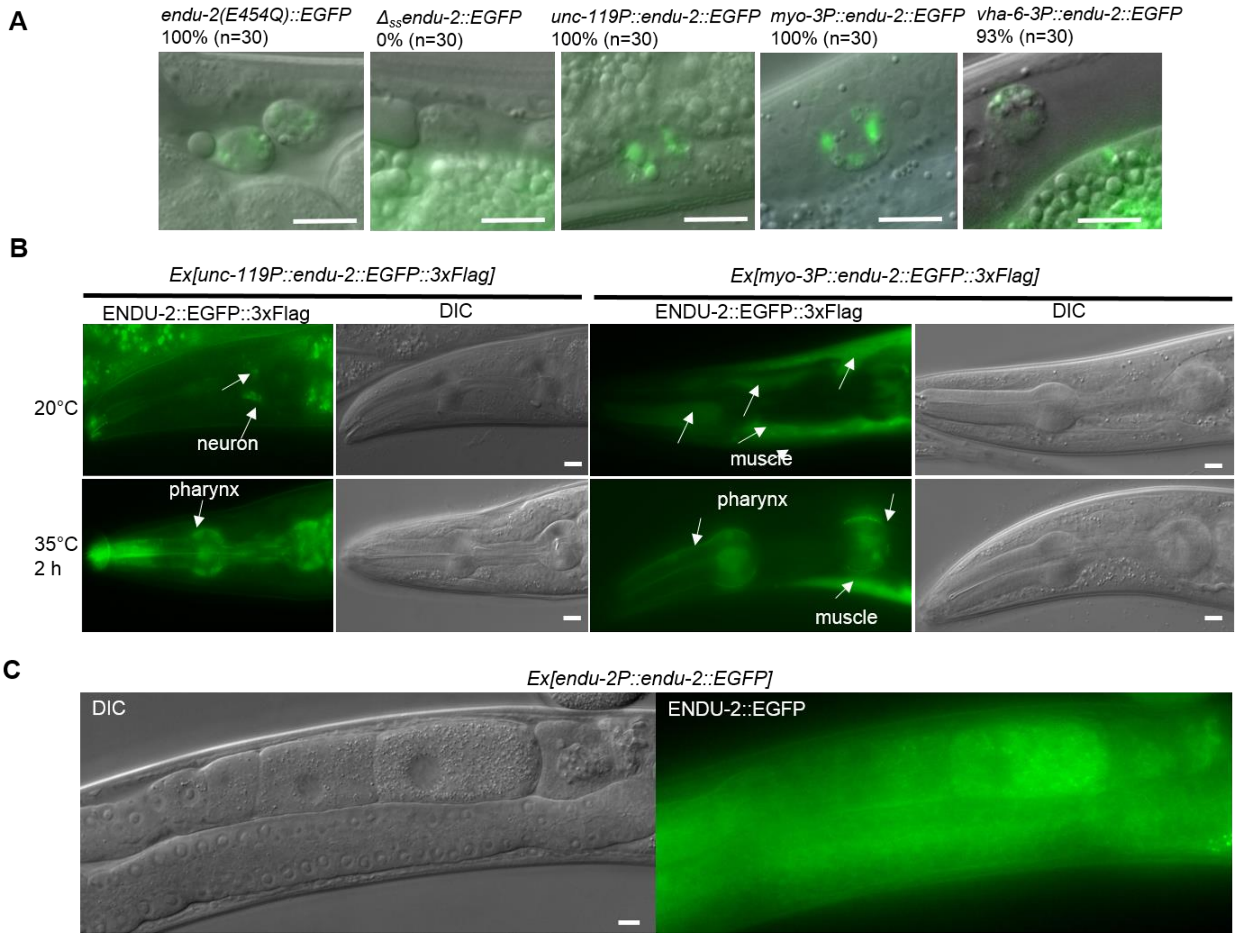


**Fig. S4 ENDU-2::EGFP is a secreted protein**

A) Accumulation of serected ENDU-2::EGFP in the coelomocytes.

B) Heat stress leads to increased pharyngeal accumulation of ENDU-2::EGFP expressed from neurons and muscle cells.

C) Extra-chromosomal *endu-2::EGFP* transgene results in weak ENDU-2::EGFP signal in the oocyte (9%, n=100).

Scale bar 10 µm. This Figure is related to the main Figure 2.

**Fig. S5**

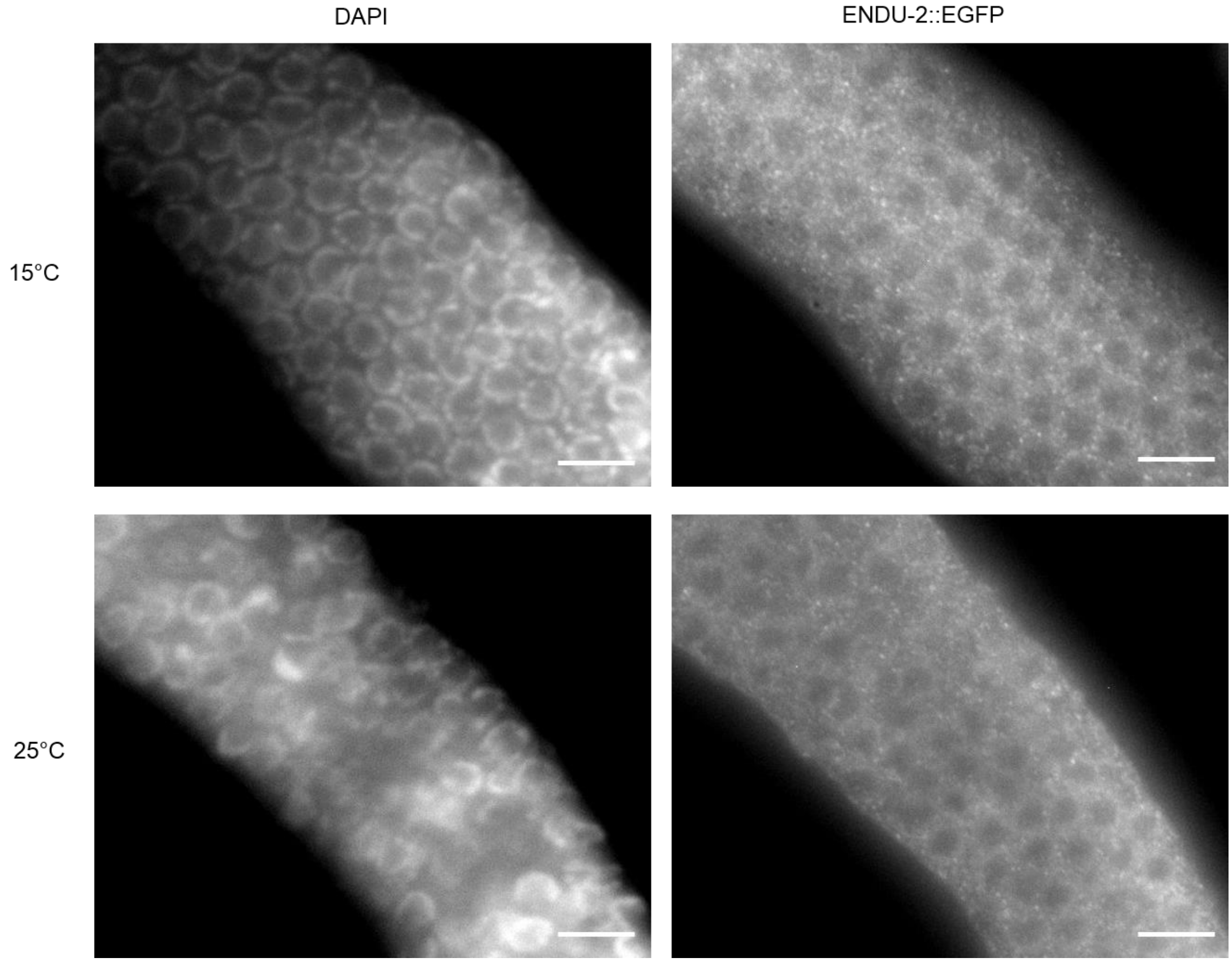


**Fig. S5. ENDU-2::EGFP is localized in the germline at both 15°C and 25°C.**

ENDU-2::EGFP in *endu-2(by190[endu-2::EGFP])* is detected with GFP antibody. Scale bar 10 µm. N=3 replicates

This Figure is related to the main Figure 2.

**Fig. S6**

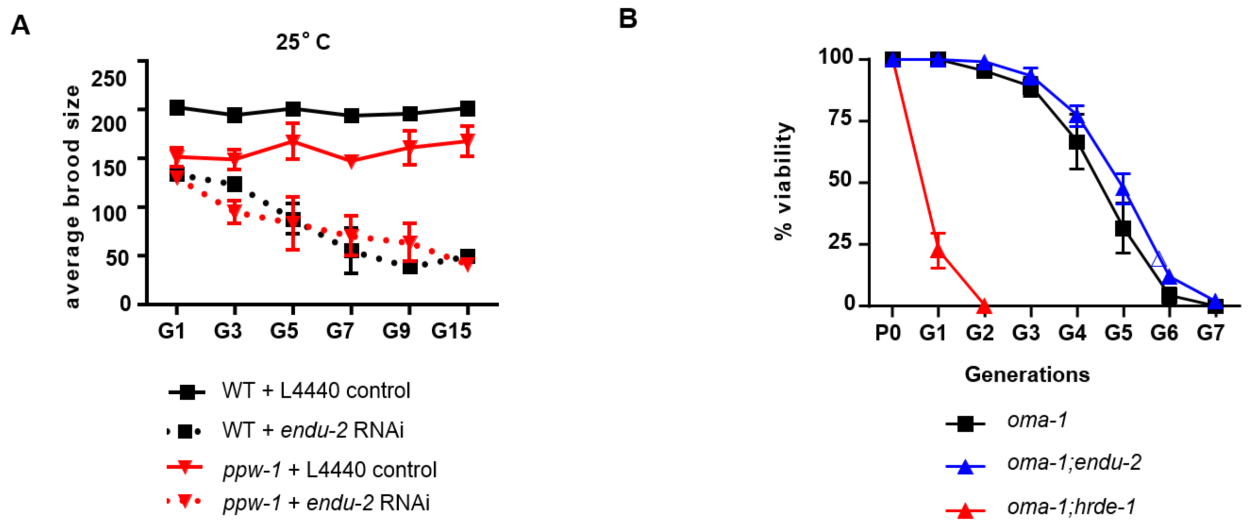


**Fig. S6 Somatic produced ENDU-2 affects reproduction without affecting multigenerational inheritance of *oma-1* RNAi in the germline.**

A) *endu-2* RNAi causes gradually reduced brood size in both wild type and *ppw-1(pk1425)* animals.

B) *endu-2(tm4977)* mutant animals are not defective in multigenerational inheritance of *oma-1* RNAi in the germline.

This Figure is related to the main Figure 3.

### Figure S7

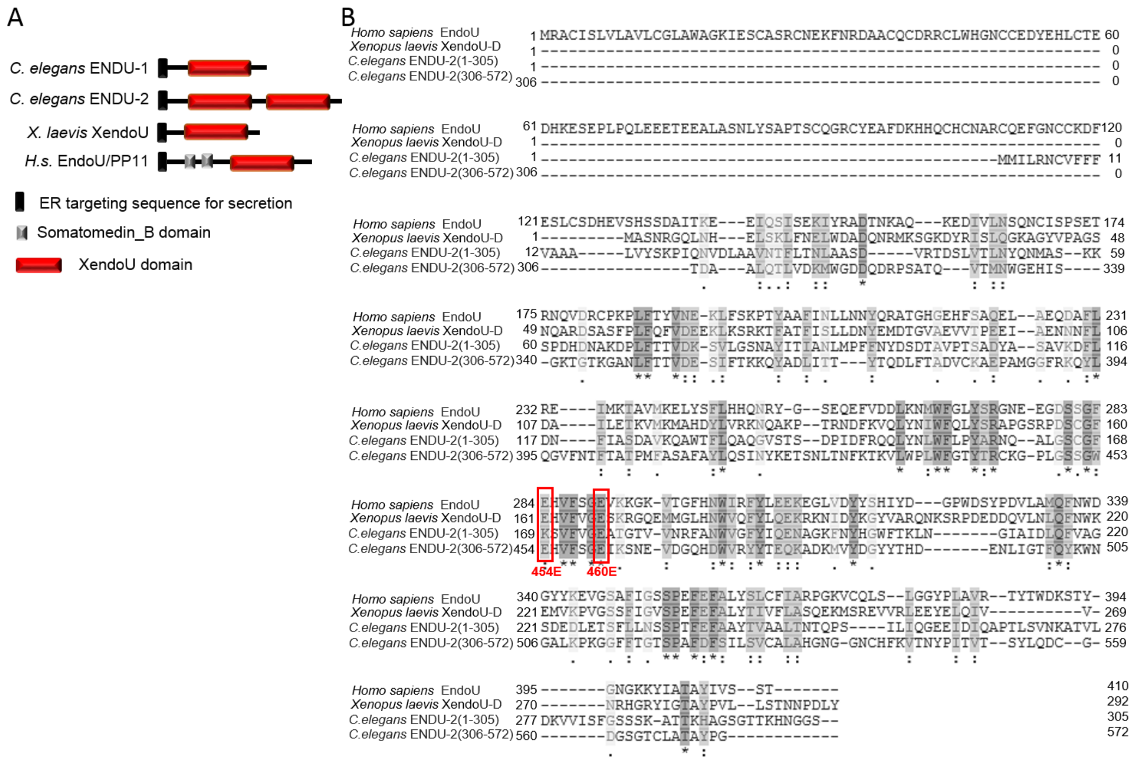


**Fig. S7 ENDU-2 is a conserved poly-U specific endoribonuclease**

A) Protein domain prediction of EndoU in *C. elegans*, *X. laevis* and *H. sapiens*.

B) Sequence alignment of human EndoU, *Xenopus* XendoU and *C. elegans* ENDU-2 suggests sequence conservation of the XendoU domains from worm to human. As *C. elegans* ENDU-2 has two XendoU domains, the alignment was performed with split protein fragments containing one of the XendoU domains (1-305 and 306-572 amino acids) each.

This Figure is related to the main Figure 4.

**Figure S8.**

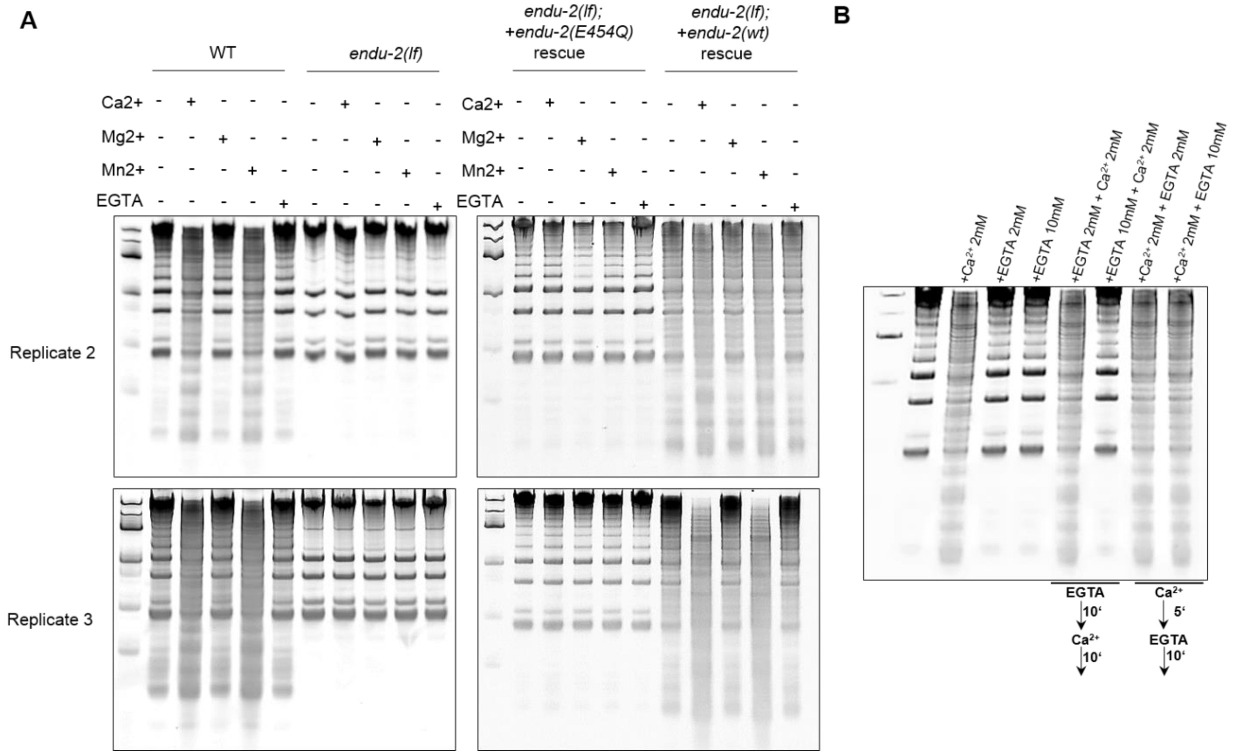


**Fig. S8 $Ca^{2+}$ and $Mn^{2+}$ promotes ENDU-2 mediated RNA decay.**

A) Two additional biological replicates showing ENDU-2 mediated bulk RNA decay in the presence of $Ca^{2+}$ and $Mn^{2+}$.

B) Incubation with $Ca^{2+}$ results in decay of total RNA isolated from wild type animals. This effect is blocked by pre-incubation with EGTA.

This Figure is related to the main Figure 4.

**Figure S9.**

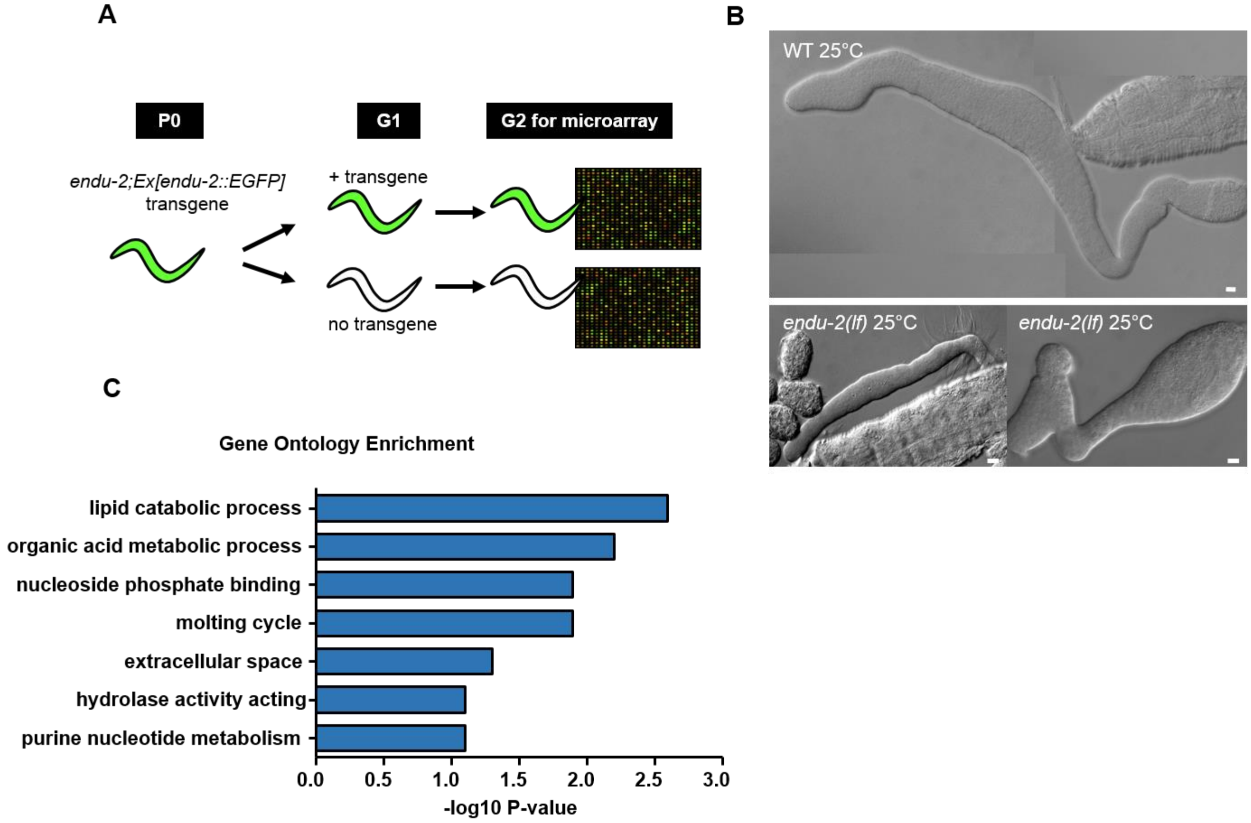


**Fig. S9 Transcriptomic analysis with microarray**

A) Schematic diagram of strain preparation for microarray. The G2 granddaughter generation from one single *endu-2(lf);Ex[endu-2::EGFP]* was used for transcriptome analysis.

B) Gonad of *endu-2(lf)* mutants are smaller than wild type animals at 25°C. Scale bar 10 µm.

C) GO term analysis of the ENDU-2 targets in the soma.

This Figure is related to the main Figure 5.

**Figure S10.**

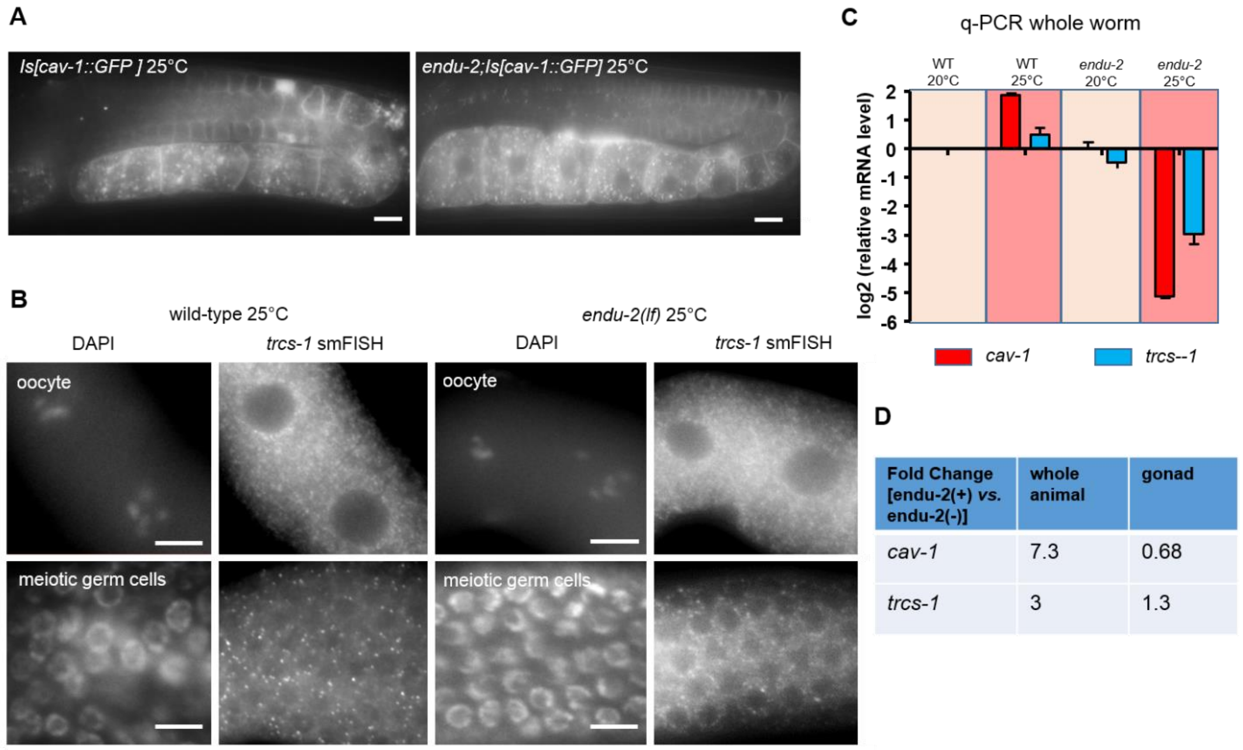


**Fig. S10. Comparison of different assays to determine expression level of *cav-1* and *trcs-1* in germline.**

A) ENDU-2 does not significantly affect the expression level of *cav-1::GFP* in the germline.

B) smFISH staining reveals no significant fold change of *trcs-1* mRNA in the germline upon *endu-2* knock-down.

C) q-PCR results suggest strongly decreased mRNA level of both *trcs-1* and *cav-1* in the absence of *endu-2* at elevated temperature.

D) Transcriptomic studies using whole animal or gonadal extracted RNA vary in their *cav-1* and *trcs-1* expression levels.

Scale bar 10 µm. This Figure is related to the main Figure 5.
